# Increased resolution in the face of conflict: phylogenomics of the Neotropical bellflowers (Campanulaceae: Lobelioideae), a rapid plant radiation

**DOI:** 10.1101/2022.01.09.475565

**Authors:** Laura P. Lagomarsino, Lauren Frankel, Simon Uribe-Convers, Alexandre Antonelli, Nathan Muchhala

**Affiliations:** Shirley C. Tucker Herbarium, Department of Biological Sciences, Louisiana State University, Baton Rouge, Louisiana, USA; Department of Biology, University of Missouri-St. Louis, St. Louis, Missouri, USA; Department of Botany, University of Wisconsin-Madison, Madison, Wisconsin, USA; Invitae Corporation, San Francisco, California, USA; Royal Botanic Gardens, Kew, TW9 3AE, UK; Gothenburg Global Biodiversity Centre, Department of Biological and Environmental Sciences, University of Gothenburg, Gothenburg, 405 30, Sweden; Department of Plant Science, University of Oxford, Oxford, UK

**Keywords:** *Burmeistera*, Campanulaceae, *Centropogon*, convergent evolution, cytonuclear discordance, gene tree conflict, Lobelioideae, museomics, Neotropics, phylogenomics, rapid radiation, *Siphocampylus*, taxonomy

## Abstract

**Background and Aims:** The centropogonid clade (Lobelioideae: Campanulaceae) is an Andean-centered rapid radiation characterized by repeated convergent evolution of morphological traits, including fruit type and pollination syndromes. While previous studies have resolved relationships of lineages with fleshy fruits into subclades, relationships among capsular species remain unresolved, particularly along the phylogenetic backbone. This lack of resolution has impeded reclassification of non-monophyletic genera, whose current taxonomy relies heavily on traits that have evolved multiple times within the clade.

**Methods:** Targeted sequence capture using a probeset recently developed for the centropogonid clade was used to obtain phylogenomic data from DNA extracted from both silica-dried and herbarium leaf tissue. These data were used to infer relationships among species using concatenated and partitioned species tree methods, as well as to quantify gene tree discordance.

**Key Results:** While silica-dried leaf tissue resulted in generally more and longer sequence data, the inclusion of herbarium samples improved phylogenetic reconstruction. Relationships among baccate lineages are similar to those inferred by previous studies, though they differ within and among capsular lineages. We improve resolution of *Siphocampylus*, which forms ten groups of closely related species to which we provide informal names that largely do not correspond to current infrageneric taxonomy. Two subclades of *Siphocampylus* and two individual species are rogue taxa whose placement differs widely across analyses. Gene tree discordance is high.

**Conclusions:** The first phylogenomic study of the centropogonid clade considerably improves our understanding of relationships in this rapid radiation. Differences across analyses and the possibility of additional lineage discoveries still hamper a solid and stable reclassification. Rapid morphological innovation corresponds with a high degree of phylogenomic complexity, including cytonuclear discordance, nuclear gene tree conflict, and well-supported differences between analyses based on different nuclear loci. Taken together, these results point to a potential role of hemiplasy underlying repeated convergent evolution. This hallmark of rapid radiations is likely present in many other species-rich Andean plant radiations.

## Introduction

The centropogonid clade of Neotropical bellflowers (Campanulaceae: Lobelioideae) is a species-rich, phenotypically diverse lineage of ∼550 species endemic to the Neotropics, with highest species richness in the Andean cordilleras of western South America. This clade comprises the genera *Centropogon*, *Burmeistera*, and mainland (i.e., non-Caribbean) species of *Siphocampylus*. Their high species richness and young age, with all species tracing back to ∼5 million years, make the centropogonids one of the fastest plant radiations documented to date (Lagomarsino *et al*. 2016).

The centropogonid clade is morphologically diverse. As in other rapid Andean radiations, including the iconic lupines (Hughes and Eastwood 2006), growth forms in the clade are variable, including vines with woody bases, herbs, xerophytes, shrubs, and trees. Fruits are varied; *ca.* 40% of the species produce dry capsules, while *ca.* 60% of species produce fleshy berries, which have evolved multiple times independently (Lagomarsino *et al*. 2014). However, floral morphology is arguably the most impressive aspect of the clade’s phenotypic diversity: their tubular flowers range in length from less than 1 cm to more than 10 cm, with petals in nearly all colors of the rainbow, including white, green, yellow, red, and orange. These floral forms have evolved to attract two principal classes of pollinators: hummingbirds and bats. From an ancestral hummingbird pollination syndrome, bat pollination has evolved *ca*. 13 times independently (Lagomarsino *et al*. 2017). Pollination biology within the clade is well-studied, including in the predominantly bat-pollinated genus *Burmeistera* (Muchhala 2003, 2006a; Muchhala and Potts 2007) and *Centropogon nigricans*, a species whose very long flowers are a product of a coevolutionary arms race with *Anoura fistulata*, a bat whose ribcage is modified to store its tongue, the longest among mammals relative to body size (Muchhala 2006b). Many other species are adapted to hummingbird pollination, including the eucentropogonid clade of curved flowers that are largely pollinated by the specialist sicklebill hummingbird (Stein 1992; Boehm *et al*. 2018).

The incredibly variable morphology in the centropogonid clade has been a challenge to taxonomists. Fruit type delimits genera, with *Burmeistera* and *Centropogon* producing fleshy berries and *Siphocampylus* producing dry capsules. In addition to their fleshy fruit, *Burmeistera* is further defined by its ebracteolate pedicels, isodiametric seeds, and generally inflated corolla (Lammers 1998). Subgeneric taxonomy predominantly relies on floral morphology (Wimmer 1943, 1955), often with traits directly associated with pollination syndromes. Previous phylogenetic studies, which largely relied on a small number of plastid loci and the nrDNA ITS, have shown that the current classification of the centropogonid genera has resulted in rampant non-monophyly of taxa at both the generic and subgeneric levels (Knox *et al*. 2008; Antonelli 2008; Lagomarsino *et al*. 2014), though support for the monophyly of *Burmeistera* within the centropogonid clade is strong (Bagley *et al*. 2020). The rampant non-monophyly is not surprising due to the current classification scheme’s heavy reliance on traits that have undergone convergent evolution in response to selection from biotic or abiotic selective pressures. Despite non-monophyly of taxa, there is strong support for five baccate (i.e., berry-producing) lineages (Lagomarsino *et al*. 2014), each with their own set of morphological tendencies and which correspond closely (though not exactly) to taxa previously proposed (McVaugh 1965). However, resolution among capsular lineages (i.e., *Siphocampylus*) remains poor.

This poor resolution is likely due to the fact that the centropogonid clade is a rapid, recent radiation. Analyses that draw from loci from across the genome and model incomplete lineage sorting are likely to be better models of its evolutionary history than the concatenation analyses of plastid loci of previous analyses. Fortunately, the last decade has seen an increase in genome-scale datasets in diverse subfields of biology as high-throughput sequencing has become cheaper and more efficient (McKain *et al*. 2018). This has resulted in a profound change in the methodology of phylogenetics (Edwards 2009; Bravo *et al*. 2019; Rothfels 2021). As genomic-scale datasets become increasingly easy to generate, the number of loci and taxa sampled in individual phylogenetic studies is increasing. This has allowed the development and application of evolutionary models that incorporate information from the distinct evolutionary histories represented in different genes. Methods are now available that model incomplete lineage sorting (Vachaspati and Warnow 2015; Zhang *et al*. 2017; Ogilvie *et al*. 2017), polyploidy (Jones *et al*. 2013), and introgression (Than *et al*. 2008; Solís-Lemus *et al*. 2017) into phylogenetic inference. In addition to taking advantage of the signal of diverse genes from the nuclear genome, deeper taxon sampling is now feasible due to the dual effect of decreasing costs of DNA sequencing and the development of methods that effectively isolate putatively homologous sequence regions from diverse taxa (McKain *et al*. 2018). It is uncontroversial to say that we are in the midst of a phylogenetic revolution.

This data revolution has led to an increased importance of genetic data from historical specimens (Lendemer *et al*. 2020). While the degraded DNA of their specimens was a major impediment to their inclusion in molecular phylogenies in the era of Sanger sequencing, natural history collections are now a major source of molecular phylogenetic data. Specimens from natural history collections, including herbaria, are commonly used in phylogenomic analyses (Bakker 2017; Brewer *et al*. 2019; Alsos *et al*. 2020; Kates *et al*. 2021). Many of these studies use hybrid enriched targeted sequence capture, a cost-effective method that is robust to low-quality, low quantity DNA due to the high specificity of the probes used to isolate targeted genomic regions which uses RNA baits to isolate a targeted subset of the genome (Faircloth *et al*. 2012; Lemmon *et al*. 2012; Weitemier *et al*. 2014)

In this manuscript, we highlight the utility of sequence capture in incorporating historical specimens in the phylogenomic analysis of the centropogonid clade of Neotropical bellflowers. Using targeted sequence capture of loci recently developed for its subclade *Burmeistera* (Bagley *et al*. 2020), we infer the most data-rich phylogeny of the entire centropogonid clade to date, inferred with both species tree and concatenation methods. Despite a high degree of gene tree discordance and differences across separate analyses, we confirm the monophyly of previously named berry-producing lineages, and further identify allied groups of *Siphocampylus* species. We end with a discussion of the potential sources of gene tree discordance in the centropogonid clade, its implications for morphological evolution, and the continued difficulty of resolving phylogenies of rapid radiations, even in the face of large phylogenomic datasets.

## Materials and Methods

### Taxon Sampling

We sampled 152 species from five genera of Lobelioideae. 108 of these were from field collected tissue and 44 from herbarium specimens. Taxon sampling was densest in *Burmeistera* (74 spp., or 49% of the sampling effort). A full taxon list and voucher information can be found in Appendix S1.

There were two goals in sampling design: to have dense phylogenetic sampling within *Burmeistera* for species-level phylogenetics (Bagley *et al*. 2020) and broad representative sampling for *Centropogon* and *Siphocampylus* to resolve the phylogeny of the clade as a whole and facilitate future reclassification efforts. The broad-scale taxon sampling used the phylogeny of Lagomarsino *et al*. (2014) as a reference from which to include a full set of known subclades within the centropogonids. Species that represented the phylogenetic breadth of the centropogonid clade and its closest relatives were sampled from both field-collected tissue and herbarium collections. Field collected tissues primarily came from previous work by AA, LPL and NM. The herbarium at the Missouri Botanical Garden (MO; code following Index Herbariorum, Thiers et al 2013 onwards) was the primary source of herbarium tissue. This herbarium was most appropriate for this study due to its extensive collection of Neotropical specimens, including collections made by a taxonomic expert in *Centropogon*, Dr. Bruce A. Stein, from the mid to late 1980s. These specimens are both expertly identified and of an appropriate age to make effective DNA extraction likely. Across our sampled herbarium specimens, the range of ages at time of extraction was 4–77 years, with a mean age of 15.3 years. Multiple species that have never before been sampled in a phylogenetic analysis were included, especially from the polyphyletic genus *Siphocampylus*.

### Molecular lab work and sequence generation

DNA extraction followed a modified CTAB protocol (Doyle and Doyle 1987). Approximately 100 mg of leaf tissue was used in these extractions for both herbarium specimens and field-collected tissue; due to tissue availability, a smaller amount of leaf tissue was used for a limited number of herbarium samples. No special treatment was given to tissue from herbarium specimens relative to silica-dried tissue. DNA concentration was assessed using a Qubit Fluorometer, followed by agarose gel electrophoresis to determine whether or not high molecular weight DNA was present.

Sequence capture targeted a set of 745 nuclear loci that were developed from transcriptomes for phylogenetic utility within *Burmeistera*, a subclade of Neotropical bellflowers (see (Bagley *et al*. 2020) for details). Illumina library preparation, target enrichment sequence capture, and sequencing were outsourced to RAPiD Genomics LLC (Gainesville, FL, USA). Degraded DNA, as inferred via gel imaging and common in our herbarium samples, was not sheared during library preparation. Sequencing was performed on an Illumina HiSeq 3000 (Illumina Inc., San Diego, CA, USA), generating 150 bp paired-end reads.

### Data processing, assembly, and extraction into targeted loci

phyluce v1.5 (Faircloth 2016) was used to process raw data into gene trees. This pipeline differs from other common sequence capture assemblers used in plants (e.g., HybPiper (Johnson *et al*. 2016) in that contigs are first assembled *de novo* and then mapped onto the target sequences. We first used illumiprocessor v2.0.8, a Trimmomatic wrapper built into phyluce (Faircloth 2013; Bolger *et al*. 2014), to trim adapter contamination and low quality bases from raw reads under default settings. We then assembled the clean reads into contigs within phyluce using the default settings of Trinity v2.6.6 (Grabherr *et al*. 2011). The newly assembled contigs were then matched to probes with 80% minimum coverage and 80% minimum identity required for a match.

Individual loci extracted with phyluce were aligned and edge trimmed with MAFFT v7.130b (Katoh and Standley 2013) under default settings. Gblocks v0.91b (Castresana 2000) was used to internally trim poorly aligned regions under default settings.

To compare the assembly of DNA from herbarium specimens and silica-collected tissue, we calculated basic summary statistics of the phyluce assemblies between these two tissue types (e.g., average number of assembled basepairs, number of assembled contigs, average contig length, and number of long loci [e.g., >1kb bp]).

### Gene tree inference and curation

Individual gene trees were inferred in RAxML v8.2.11 using rapid bootstrapping with a GTR model of molecular evolution and between 78-130 bootstrap replicates. After exploratory species tree analyses, we filtered our data to improve phylogenetic inference (Molloy and Warnow 2018). Using contig length as a proxy, we first filtered out taxa with low copy data assemblies (i.e., all contigs less than 1000 bp). These 13 taxa (from an initial 152) were primarily from DNA extracted from herbarium specimens with low quantity of input DNA (Appendix S2). Second, we filtered individual loci: we removed loci that had less than 75% representation of the remaining 139 taxa, as well as loci that were shorter than 500 bp.These filtering steps left 96 loci for 139 taxa, which were used in all downstream analyses.

### Phylogenomic Analysis

To infer phylogenetic relationships, we performed a concatenated analysis on a combined, partitioned alignment of loci in RAxML using rapid bootstrapping. PartitionFinder2 (Lanfear *et al*. 2017) was used to determine the optimal partitioning scheme of loci. Subsequently, three different methods were then utilized to estimate coalescent-based species trees from: ASTRAL-III v5.6.2, (Zhang *et al*. 2017), ASTRID v1.4 (Vachaspati and Warnow 2015) using the BioNJ method, and SVDQuartets implemented in PAUP v4.0a162 (Swofford 2001; Chifman and Kubatko 2014). ASTRAL-III and ASTRID used the individual gene trees as input; a concatenated alignment of all filtered loci was used to estimate the species tree with 100 bootstrap replicates in SVDQuartets.

We assessed gene tree discordance by mapping rooted gene trees to the RAxML topology using PhyParts (Smith *et al*. 2015) and subsequently visualized results using PhyPartsPieCharts. (https://github.com/mossmatters/phyloscripts/tree/master/phypartspiecharts).

### Comparison to previous analyses

To explore differences with past studies, we constructed a tanglegram of relationships among species included in both our RAxML concatenated phylogeny and the plastid phylogeny from Lagomarsino et al. (2014) using the tanglegram function of the dendextend R package (Galili 2015).

## Results

### Taxon sampling and target capture success

Sequence data for targeted loci were obtained for all included taxa, though the quantity was variable, ranging from only 6605bp in *Burmeistera breviflora* to 502,363bp in *Lysipomia pumila*. The number of contigs recovered ranged from 28 to 669, with an average of 343. Median length of these ranged from 222 to 933bp (with an average median length of 646bp). Our final dataset included 139 accessions, representing 68 species of *Burmeistera*, 26 species of *Centropogon*, 42 species of *Siphocampylus*, *Lysipomia pumila,* and *Lobelia tupa*. A complete list of target capture summary statistics can be found in Appendix S2.

### Performance of herbarium vs. silica-dried tissue

Across all of our samples, silica-dried tissue outperformed herbarium tissue in most summary statistics: the mean total assembled base pairs for silica tissue was ca. 332kp (vs. 181 kp for herbarium specimens; t-test p-value <0.0001); the mean average contig length was 951 bp for silica-dried tissue (vs. 525 bp; t-test p-value <0.0001); and the mean number of contigs greater than 1kp was 83 in silica-dried tissue (vs. 30 from herbarium specimens; t-test p-value <0.0001) (Fig. 2). However, herbarium and silica-dried tissue did not statistically significantly differ in the total number of contigs recovered (331 vs 351 contigs; t-test p-value=0.39).

Based on preliminary analyses of our data, we considered any taxon lacking any contigs greater than 1000 bp as insufficient quality for phylogenomic analysis. In preliminary phylogenetic analyses, these samples were attracted to the root of the phylogeny, forming grades successively sister to larger clades, a known artifact in phylogenomic analyses (Hosner *et al*. 2016). With this criterion, all field-collected specimens had enough data of sufficient quality to be included in phylogenetic analysis, while only 31 samples (representing 70.5% of the total) herbarium-collected specimens could be included. Of those that failed, a majority (11/13) had less than 250 ug of input DNA. However, one sample that failed had more than 1300 ug of input DNA (i.e., *C. tessmannii*), and at the other extreme, our sample with the smallest amount of input DNA (i.e., *B. curviandra*) had efficient enough capture to be included.

### Broad phylogenetic patterns

Building on recent results within *Burmeistera* (Bagley *et al*. 2020), we expand upon previous understanding of relationships within the broad centropogonid clade relative to previous analyses, the majority of which have relied on primarily or entirely plastid sequence (Knox *et al*. 2008; Antonelli 2009; Lagomarsino *et al*. 2014; Bagley *et al*. 2020). This is the first nuclear phylogenomic study that targets the entire centropogonid clade. Phylogenetic relationships of *Centropogon* and *Siphocampylus* species are presented in Fig. 2-3, while phylogenies that include *Burmeistera* are depicted in Figures S1-4. PhyParts results revealed a high degree of gene tree discordance in our dataset: most gene trees had conflicts other than the most common gene tree-species conflict at the majority of nodes (indicated as red portion in the pie charts in Fig.2, Fig. S5). A review of individual gene trees suggests that there are many conflicting phylogenies, not a small number of conflicts that are repeated throughout our dataset.

Despite this, our results significantly advance our understanding of the relationships within and between subclades. At a broad scale, we identify two major clades of centropogonids that are supported in all our analyses: one that includes the burmeisterid, brevilimbatid, peruvianid, and colombianid *Centropogon* species in addition to various lineages of *Siphocampylus*; and a second that includes the eucentropogonid clade and various lineages of *Siphocampylus*. In addition, two subclades and three individual species of *Siphocampylus* species have inconsistent placement across our analyses.

The last comprehensive phylogenetic analysis of all three centropogonid genera (Lagomarsino *et al*. 2014) resolved the relationships among the berry-producing lineages (i.e., those taxa currently classified as *Centropogon* and *Burmeistera*) into five well-support subclades (i.e., burmeisterids, the brevilimbatids, the peruvianids, the colombianids, and the eucentropogonids) using plastid sequences. We find strong support for the monophyly of four of those subclades (i.e., the burmeisterids, brevilimbatids, peruvianids, and eucentropogonids) in all of our analyses. The *Centropogon* species of the remaining named subclade, the colombianids, are closely related in most of our analyses (i.e., monophyletic in ASTRID, SVDquartets, RAxML; Fig. 2, 3BC and paraphyletic in ASTRAL; Fig. 3A); however, the unstable placement of *S. nematosepalus*, currently considered to belong to this clade, renders it non-monophyletic in every analysis. *Siphocampylus nematosepalus* is an outlier within the colombianid clade for its capsular fruit, which is hypothesized to represent a reversal from baccate ancestors (Lagomarsino *et al*. 2014).

While we largely confirm phylogenetic relationships within berry-producing taxa (i.e., *Burmeistera* and *Centropogon*), as outlined above, our results provide novel insights into the relationships of capsular taxa (i.e., “*Siphocampylus*”), which were largely unresolved in (Lagomarsino *et al*. 2014). As in previous studies, we find strong support for the polyphyly of species currently classified in *Siphocampylus*, and we identify ten groups of *Siphocampylus* species that are consistently resolved as closely related. We have placed new informal names on these *Siphocampylus* lineages (Fig. 2), and include a description of them below in the context of the broader clades in which they occur.

### Clade 1

The first major clade includes at least seven distinct lineages (i.e., the burmeisterid, the brevilimbatid, the peruvianid and the colombianid *Centropogon* subclades, and the *giganteus*, *umbellatus*, and *odontosepalus* groups of *Siphocampylus* species, as well as *S. nematosepalus* and *S. smilax*). Two additional lineages of *Siphocampylus* (i.e., the *furax* and *andinus* clades) are inferred to belong in Clade 1 in the ASTRAL analysis only, and *S. corymbifer* is inferred to belong to it in all analyses except SVDquartets (Fig. 2, 3). While the membership of this clade differs across analyses based on the placement of these taxa, it is well-supported in most analyses (ASTRAL/ ASTRID/ SVDQuartets/ RAxML support: 0.96/0.68/92/99).

The burmeisterid subclade, which comprises *Burmeistera* and various bat pollinated *Centropogon* species that share aspects of their gross floral morphology with *Burmeistera* (represented by *C. nigricans* and *C. smithii* here), is inferred as monophyletic in all analyses (0.55/0.85/100/67). Within the burmeisterid subclade, the monophyly of *Burmeistera*, represented by 68 taxa in our sampling, is moderately- to well-supported (0.7/0.99/95/78). The well-supported brevilimbatid subclade (1/0.99/100/71) is sister to the burmeisterid subclade in all analyses, consistent with results from (Lagomarsino *et al*. 2014). This subclade comprises species of *Centropogon* with orange to red flowers generally with yellow corolla limbs and is represented by four taxa in our sampling (i.e., *C. macbridei*, *C. grandidentatus*, *C. erythraeus*, and *C. aslcepiadeus*). Three robust, green-flowered species that we refer to as the *giganteus* grade are successively sister to the brevilimbatids + burmeisterids; together, these three groups have moderate support (0.71/0.72/63/72). The relationship between the burmeisterid, brevlimibatid, and giganteus grade is consistent across analyses, and further consistent with Lagomarsino et al (2014).

Three remaining major groups (i.e., the peruvianid *Centropogon* clade and the *umbellatus* and *odontosepalus* groups of *Siphocampylus*) and two species (i.e., *S. smilax* and *S. corymbifer*) are always included in Clade 1; however, their relationship differs across analyses. In both the ASTRAL and ASTRID analyses, these groups form basal polytomies, with resolution within individual and between individual subclades.

Across analyses, we find evidence for a close relationship between peruvianid *Centropogon* subclade and the *umbellatus* group of *Siphocampylus*, both groups of generally green-to-white flowered species that are either sister to, are successively sister to, or form a polytomy with the burmeisterid-giganteus clade. The peruvianids, a subclade of robust shrubs with large red or white flowers that is restricted to the Central Andes of Peru and Bolivia and represented by nine tips in our sampling (*C. dianae*, *C. simulans*, *C. perlongus*, *C. viriduliflora*, *C. mandonis*, *C. brittonianus*, and three accessions of *C. incanus*), is strongly supported as monophyletic in all analyses (1/1/100/100). The *umbellatus* group, represented by five species in our sampling, is phenotypically similar to the peruvianid subclade, though these species produce capsules and not berries; these robust shrubs include species with both typically hummingbird (e.g., *S. boliviensis*) and bat (e.g., *S. tunicatus*) pollinated flowers (Lagomarsino and Santamaría-Aguilar 2016). The relationships between species in the *umbellatus* group differ across analyses; it is monophyletic in the ASTRAL analysis (0.74) and a grade or unresolved in the remainder. Together, these two major groups form a clade (i.e., ASTRID and RAxML [the latter with the exception of *S. tunarensis* and *S. boliviensis*]) or grade (i.e., ASTRAL, SVDquartets).

We additionally find strong support for a close relationship between the colombianid subclade of *Centropogon* and a group of *Siphocampylus* species that we refer to as the *odontosepalus* group. Together, these species are strongly supported as monophyletic across all analyses (0.91/0.99/96/100). These phenotypically similar groups are both generally suffrescent subshrubs with narrow, bright pink flowers found in the northern Neotropics; the colombianid subclade is distributed from southern Central America to northern South America, while the *odontosepalus* group comprises *Siphocampylus* species from northern South America (i.e., *S. odontosepalus* from Venezuela*, S. longibracteolatus* from Colombia, and *S. planchonis* from Colombia and Venezuela). The relationship between these groups differs across analyses; they are either sister clades (RAxML; Fig. 2), or the colombianid species form a grade to a monophyletic *odontosepalus* group (ASTRAL; Fig. 3A), or the *odontosepalus* species form a grade to a monophyletic colombianid group (ASTRID, SVDQuartets; Fig. 3B-C). In the SVDquartets and RAxML analyses, these groups are sister to the clade that includes both the major of Clade 1 species (i.e., the clade defined by the common ancestor of the burmeisterids and the *umbellatus* group). As previously noted, the monophyly of the colombianid clade *sensu* (Lagomarsino *et al*. 2014) is not supported due to the unstable placement of *S. nematosepalus*, a capsular species from montane Costa Rica (see discussion of rogue taxa below).

Finally, *S. smilax* is either sister to the remainder of Clade 1 or forms a part of a basal polytomy with its other major subclades (Fig. 2, 3). This is a green-flowered species from Bolivia with a thickened, succulent woody base that is likely an adaptation for water storage in the relatively arid environments in which it occurs— an uncommon habitat for the centropogonid clade.

### Clade 2

The second major clade includes the eucentropogonid clade and five subclades of *Siphocampylus* species. Four of these subclades, together referred to as the *scandens* grade, are closely related in all analyses, which we collectively refer to as the *scandens* grade: i) the *correoides* subclade (0.95/1/98/100), comprising *S. correoides*, *S. ayersiae*, and *S. chloroleucus*; ii) the *nobilis* subclade (1/0.99/0.98/92), comprising *S. scandens* and *S. nobilis*; iii) the *actinothrix* subclade (0.77/1/100/100), comprising *S. actinothrix*, *S. antonelli*, and *S. vatkeanus*; and iv) *S. asplundii.* We refer to the fifth group as the *tupaeformis* clade (0.59/0.77/100/100), which corresponds to *S. fiebrigii*, *S. aureus*, *S. macropodus*, *S. werdermannii*, *S. krauseanus*, *S. obovatus*, *S. virgatus*, and *S. tupaeformis* in our sampling.

Species in the *tupaeformis* clade tend to be erect, but short-statured shrubs with relatively small, hummingbird-pollinated flowers. While each of these subclades are resolved as monophyletic in all analyses, their relationships differ across analyses. In two analyses (i.e., ASTRAL and SVDquartets), the *tupaeformis* clade is sister to the remaining groups, while it is sister to the eucentropogonid clade in the RAxML analysis (Fig. 2, 3) and forms a basal polytomy with the rest of the Clade 2 subclades in the ASTRID analysis.

The eucentropogonid clade (which corresponds to *Centropogon* subgenus *Eucentropogon*) is monophyletic with highest possible support in all analyses (1/1/100/100). This is a group of *ca*. 55 species that are often pollinated by the sicklebill hummingbird (Stein 1992; Boehm *et al*. 2018) and are characterized by extremely curved flowers with a cornute scale of fused trichomes at the apex of their ventral anthers. Relationships among eucentropogonid species are similar to those in previous analyses (Lagomarsino *et al*. 2014) and correspond to current taxonomic groups: we find that *C. cornutus* is sister to the group that includes subsections *Amplifolii* and *Campylobotrys* from (Stein 1987), and that members of *Campylobotrys* are monophyletic. Only a single species of subsection *Amplifolii*, *C. congestus*, is included in our sampling, thus we cannot evaluate its monophyly. However, there is one important difference from previous analyses: we find that a currently undescribed *Centropogon* species (whose morphology suggests that it belongs to the eucentropogonid clade as it shares the synapomorphic cornute scale and curved corolla) is sister to the rest of this clade.

While the subclades of the *scandens* grade lack resolution, relationships are consistent in the two analyses in which they do not form a polytomy (i.e., ASTRAL and RAxML analyses). In these analyses, there is strong support for a sister relationship between the *correoides* and *nobilis* subclades (1.0 and 100, respectively). All members of these clades are soft-woody vines often with coriaceous leaves, though floral morphology is notably variable, from typically hummingbird-pollinated flowers that are small, narrow, pink, and relatively long (e.g., *S. scandens*) to typically bat-pollinated flowers that are large, wide, green, and relatively short (e.g., *S. ayersiae*). The two additional subclades (i.e., the *actinothrix* subclade and *S. asplundii*) form a clade in the ASTRAL and RAxML analyses (0.77 and 100, respectively), and are sister the remaining members of the *scandens* grade and the eucentropogonid clade (ASTRAL) or the eucentropogonid + tupaeformis subclades (RAxML). The *actinothrix* subclade is composed of erect woody shrubs with a combination of flowers with hummingbird (e.g., *S. antonellii*) and bat (e.g., *S. actinothrix*) pollination syndromes (Lagomarsino and Santamaría-Aguilar 2016); *S. asplundii* is a scandent shrub with a narrow, pink flower.

### Taxa of uncertain placement

The membership of Clade 1 and Clade 2 is consistent across analyses with the exception of two subclades that are resolved in very different regions of the phylogeny across analyses: i) the *furax* clade (0.98/0.99/99/100), comprising *S. furax*, *S. affinis,* and *S. brevicalyx* and ii) the *andinus* clade (1/0.99/99/100), comprising *S. imbricatus*, *S. orbignianus*, *S. rusbyanus*, *S. andinus*, *S. verticillatus*, *S. lycioides*, and *S. longipedunculatus*.

These two subclades tend to be closely related across analyses, forming a clade (ASTRAL; Fig. 3A) or grade (ASTRID, RAxML; Fig. 2, 3B), though they form part of a basal polytomy in the SVDquartets analysis (Fig. 3C). Their placement within the broader phylogeny differs across analyses. They are nested within Clade 1 in the ASTRAL analysis, are successively sister to Clade 2 in the ASTRID and RAxML analyses, and are part of a polytomy at the base of the centropogonid clade in the SVDquartets analysis.

*Siphocampylus corymbifer* is a scandent shrub with a corymb inflorescence that is atypical within the centropogonid clade and a similarly atypical widespread distribution that stretches from the Andean forests of Peru and Bolivia to the Atlantic Forest of Brazil. Its placement differs widely across analyses: ASTRAL and ASTRID place it in a polytomy at the base of Clade 1, SVDquartets places it in a polytomy at the base of the centropogonid clade, and RAxML places it sister to two other Central Andean species ( *S. tunarensis* and *S. boliviensis*) within Clade 1. The second rogue species, *S. nematosepalus*, which was previously included in the colombianid clade (Lagomarsino et al. 2014), always belongs to Clade 1 but is placed differently across analyses: ASTRAL and RAxML place it sister to the *odontosepalus* group and colombianid clade with relatively strong support (0.91 and 100, respectively), ASTRID and SVDquartets infer to be sister to the burmeisterid-brevilimbatid-giganteus groups (with support of 0.72 and 92, respectively).

## Discussion

Building on recent progress within *Burmeistera* (Bagley *et al*. 2020), we present the first phylogenomic analysis of the entire centropogonid clade. Targeted sequence capture permitted herbarium specimens to be a major source of DNA, and despite the degraded quality of herbarium DNA, its inclusion did not present major barriers. After meeting certain minimum requirements for DNA quality, sequence capture was efficient for herbarium tissue, although it underperformed compared to field-collected tissues. This study presents the most data-rich phylogeny of this group to-date, and is among the first nuclear phylogenomic studies of a species-rich cloud forest-centered Andean clade (Nevado *et al*. 2016; Vargas *et al*. 2017; Pouchon *et al*. 2018; Bagley *et al*. 2020; Acha *et al*. 2021). Inferred relationships represent significant departure from previous knowledge of phylogeny in the clade, particularly in increased resolution along the backbone of the phylogeny and among capsular species (currently recognized as the genus *Siphocampylus*). A high degree of gene tree discordance and complex patterns of morphological trait evolution characterize the centropogonid clade. These represent dual challenges to resolving phylogenies of rapid radiations and developing classification schemes in groups characterized by frequent morphological convergent evolution.

### Herbariomic data substantially improved phylogeny despite yielding significantly less sequence data

Overall, our results add to a growing body of literature showing that herbarium specimens are useful for increasing taxon sampling in phylogenomic analyses, although there are limitations to the use of this data source. The method that we used, sequence capture, does not require high molecular weight DNA, and instead is robust to low quantity, low quality input DNA (Hart *et al*. 2016; Brewer *et al*. 2019). Despite this, we found that sequence data derived from herbarium samples tend to be lower quality than data from silica-collected tissue. While similar numbers of contigs were recovered from both data types, those contigs tended to be shorter in herbarium specimen data, resulting in fewer long contigs (Fig. 1). Using the number of very long contigs (in our case, those greater than 1kp) as a proxy quality of data, we removed 13 (31%) of all herbarium specimens and no silica-collected tissues. However, it is likely that additional data curation would have allowed us to include these specimens in phylogenomic analysis, and this is a goal for future studies.

**Figure 1.**
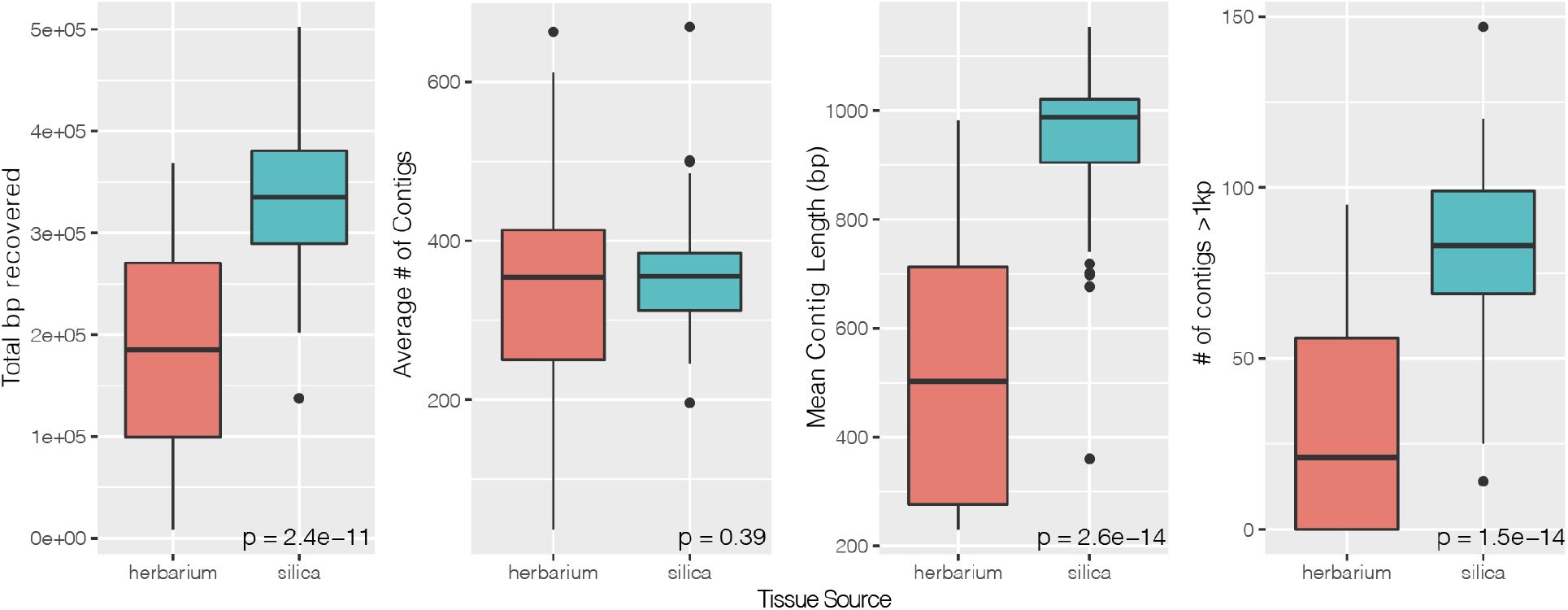
Comparison of summary statistics for targeted locus assembly for herbarium (red) vs. silica-collected (blue) leaf tissue.

Despite needing to drop many samples derived from herbarium specimens from our phylogenomic analyses, the inclusion of those that remained greatly enhanced our understanding of relationships within the centropogonid clade. For example, this approach allowed us to include species known only from a single locality (e.g., *Centropogon* sp. nov. in the eucentropogonid clade) and species endemic to countries where it is not currently feasible to collect new material (e.g., *S. odontosepalus* from Venezuela). Further, the *odontosepalus* group, which either forms a clade or a grade closely related to the colombiand clade, is newly discovered and is currently exclusively composed of samples derived from herbarium specimens; for two of their three species (i.e., *S. odontosepalus* and *S. planchonis*), our targeted sequence capture data represent the first DNA sequences generated for their species. This points to an important role of herbarium specimens in phylogenetic lineage discovery, allowing us to continue to unravel branches of the tree of life even when it would otherwise not be feasible, whether due to logistical reasons (e.g., difficulty of additional fieldwork or lack of funding) or fundamental impossibility (e.g., extinction of taxa).

Herbarium specimens will continue to be an important potential source of phylogenomic data in the years to come. Highlighting this, many recent studies include a large proportion of herbarium specimens in a sequence capture-based phylogenies (e.g., (Silva *et al*. 2017; Schneider *et al*. 2021; Thomas *et al*. 2021). However, DNA extracted from herbarium specimens is not a panacea, with data quality generally being lower when derived from herbarium specimens than from silica-dried tissue. Additionally, herbarium specimens are physical objects that are permanently damaged each time tissue is sampled (especially if no material is available in the fragment packets), and many institutions have a limit on how many times a single specimen can be destructively sampled (Rabeler *et al*. 2019). For these reasons, inclusion of herbarium specimens in phylogenomic studies should be well-justified and preferably only undertaken if no alternatives are available or if an individual specimen represents an important addition to a study (e.g., distinct morphology or locality).

### Improved phylogenetic resolution of Neotropical bellflowers

At the deepest level, we resolve two major centropogonid clades— Clade 1 and Clade 2 in Fig. 2— that are novel to this study (though two additional clades of *Siphocampylus* species, the *furax* and *andinus* clades, are variably placed within these major clades or in a basal polytomy across analyses). There are no apparent morphological synapomorphies that unite either Clade 1 or Clade 2. Both include species that span the majority of morphological variation within the centropogonid clade, including both baccate and capsular lineages; hummingbird and bat pollination syndromes; vining and erect habits; and a wide variety of pubescence types (Fig. 2).

**Figure 2.**
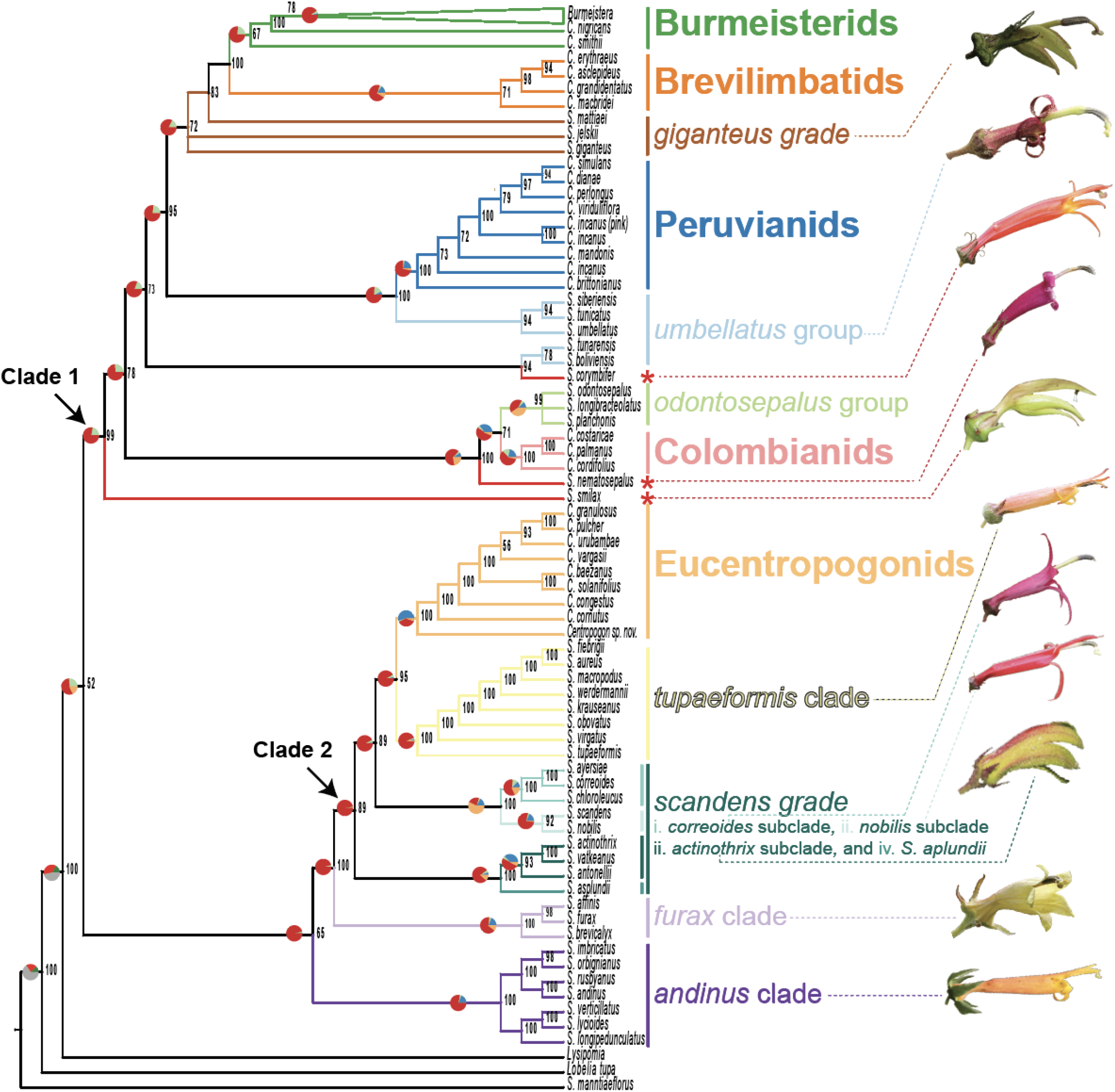
Concencated RAxML phylogeny with informal names of closely allied species presented at right, with the branches of the phylogeny corresponding to their members color-coded accordingly. Names in bolded text are from Lagomarsino et al. (2014) and correspond to the baccate lineages (i.e., species currently classified in Burmeistera and Centropogon). Names in italics correspond to capsular lineages (i.e., species currently classified in Siphocampylus). Individual rogue species are indicated by red asterisks. Numbers at nodes represent bootstrap support. Pie charts above select branches illustrate gene tree discordance as estimated in PhyParts using all gene trees along the RAxML phylogeny and visualized using PhyPartsPieCharts; colors correspond to the proportion of gene trees that fall into different categories of concordance (blue: concordant genes; green: most common conflicting bipartition; red: other conflicting bipartitions; orange: gene trees with no information). Photos of living flowers from exemplar species of the *Siphocampylu*s lineages are presented at right. From top to bottom, these are: *S. matthaei*, *S. tunarensis*, *S. corymbifer*, *S. nematosepalus*, *S. smilax*, *S. tupaeformis*, *S. correoides*, *S. scandens*, *S. actinothrix*, *S. affinis*, and *S. andinus*.

**Figure 3.**
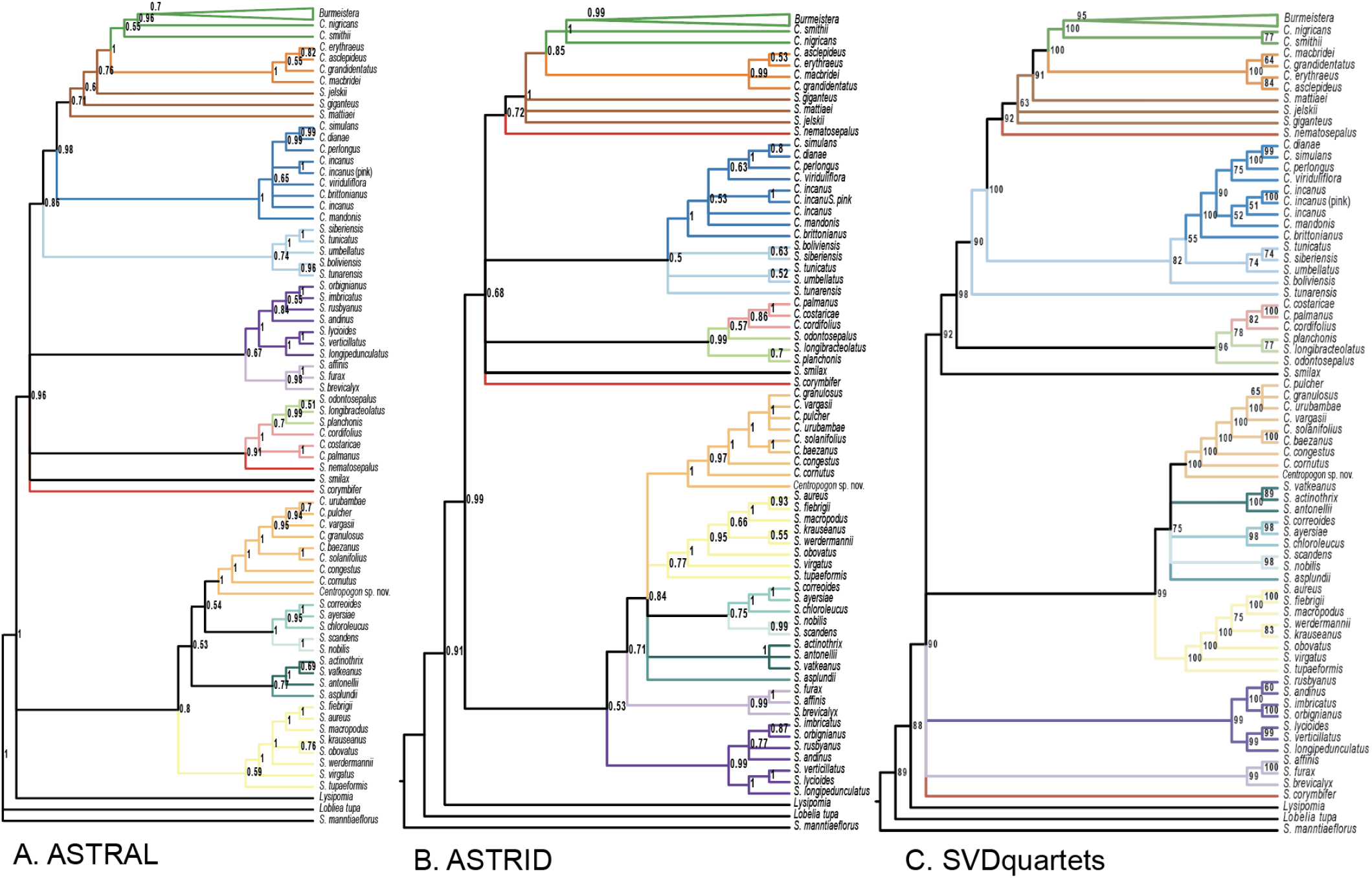
Results of species tree analyses, with consistently inferred species groups colored coded as in Fig. 2. a) ASTRAL-III analysis with local posterior probabilities values at nodes; b) ASTRID analysis with bootstrap values at nodes. C. SVDquartets phylogeny bootstrap values at nodes.

Relationships among baccate lineages are consistent across analyses and similar to previous studies. The monophyly of *Burmeistera* and four separate clades of *Centropogon* are consistently well-supported, as are the relationships between them. The fifth group of named *Centropogon* species, the colombianids (with the exception of *S. nematosepalus*), is monophyletic in all analyses except ASTRAL, and its placement within Clade 1 is consistent across phylogenies. Morphology within these baccate clades tends to be conserved beyond the fruit type that unites them, including many aspects of floral morphology, gross pollination syndrome, habit, pubescence type, and, often, biogeography (Lagomarsino *et al*. 2014).

While relationships among *Siphocampylus* have improved support relative to previous analyses, this genus, defined by its plesiomorphic capsular fruits, is far from resolved. Support, membership, and relationship between named groups of *Siphocampylus* species (as well as their relationship to baccate lineages) differs across analyses and tends to have lower support values than their baccate equivalents. Additionally, there are two rogue subclades (i.e., the *andinus* and *furax* subclades) and two rogue species of *Siphocampylus* (i.e., *S. corymbifer* and *S. nematosepalus*) that are unstable in their placement. Future research that tests for a potential hybrid origin or history of polyploidy in these lineages is likely to be fruitful. Despite these conflicts, we infer ten groups of closely allied species of *Siphocampylus* (these often forming either subclades or grades in different analyses) that are consistently identified across analyses. Contrasting with the pattern in *Centropogon* and *Burmeistera*, in which flowers tend to be similar among close relatives, subclades of *Siphocampylus* are marked by very variable floral morphology, often including multiple shifts in pollination syndromes (Fig. 2, 5).

The discrepancy between relatively high constancy of relationships in *Centropogon* and *Burmeistera* lineages as compared to *Siphocampylus* lineages is likely because a derived character state, berry fruits, defines them. Capsular fruits are the ancestral state in the centropogonid clade, from which berries have been derived on multiple occasions (Lagomarsino *et al*. 2014; Crowl *et al*. 2016). A consequence of this history is that *Centropogon* subclades are relatively young relative to both the centropogonid clade as a whole and to many individual lineages of *Siphocampylus*. Their younger age also means there has been less time for phylogenetic conflicts due to introgression and hybridization to arise. While gene tree discordance is high at the base of both baccate and capsular lineages (Fig. 2), it is notable that the nodes with the highest portion of concordant genes are at the base of baccate subclades (i.e., the eucentropogonid and peruvianid subclades).

Finally, comparing our results to the plastid phylogeny of Lagomarsino et al. (2014), we observe a pattern consistent with extensive cytonuclear discordance. Cytonuclear discordance (i.e., when the phylogenetic signal of organelle genomes– in this case, the plastome– is significantly different from phylogenetic signal of the nuclear genome) is often considered evidence of reticulate evolution (Reiseberg and Soltis 1991; Sloan *et al*. 2017), though can also be attributed to additional processes including selection and incomplete lineage sorting (Lee-Yaw *et al*. 2019).

While we find broad support for the same major baccate subclades as in the plastid phylogeny, relationships both within and between the capsular lineages differ substantially (Fig. 4). Given the extent of discordance among analyses and within nuclear gene trees, as well as the fact that the plastome represents is generally considered to be a single coalescent gene (Gonçalves *et al*. 2019, but see Doyle 2021), it is very likely that at least some of differences we observe are due to incomplete lineage sorting (Lee-Yaw *et al*. 2019; Firneno *et al*. 2020). However, much of the cytonuclear discordance we identify involves nodes deep in the phylogeny (e.g., members of the *andinus* clade of *Siphocampylus*), suggesting a potential for an ancient history of hybridization or introgression within the centropogonid clade. Given the clade’s extensive convergent evolution of morphological traits, the overlap of character states between subclades, and rapid rates of diversification, this would not be surprising and represents a promising area for future research.

**Figure 4.**
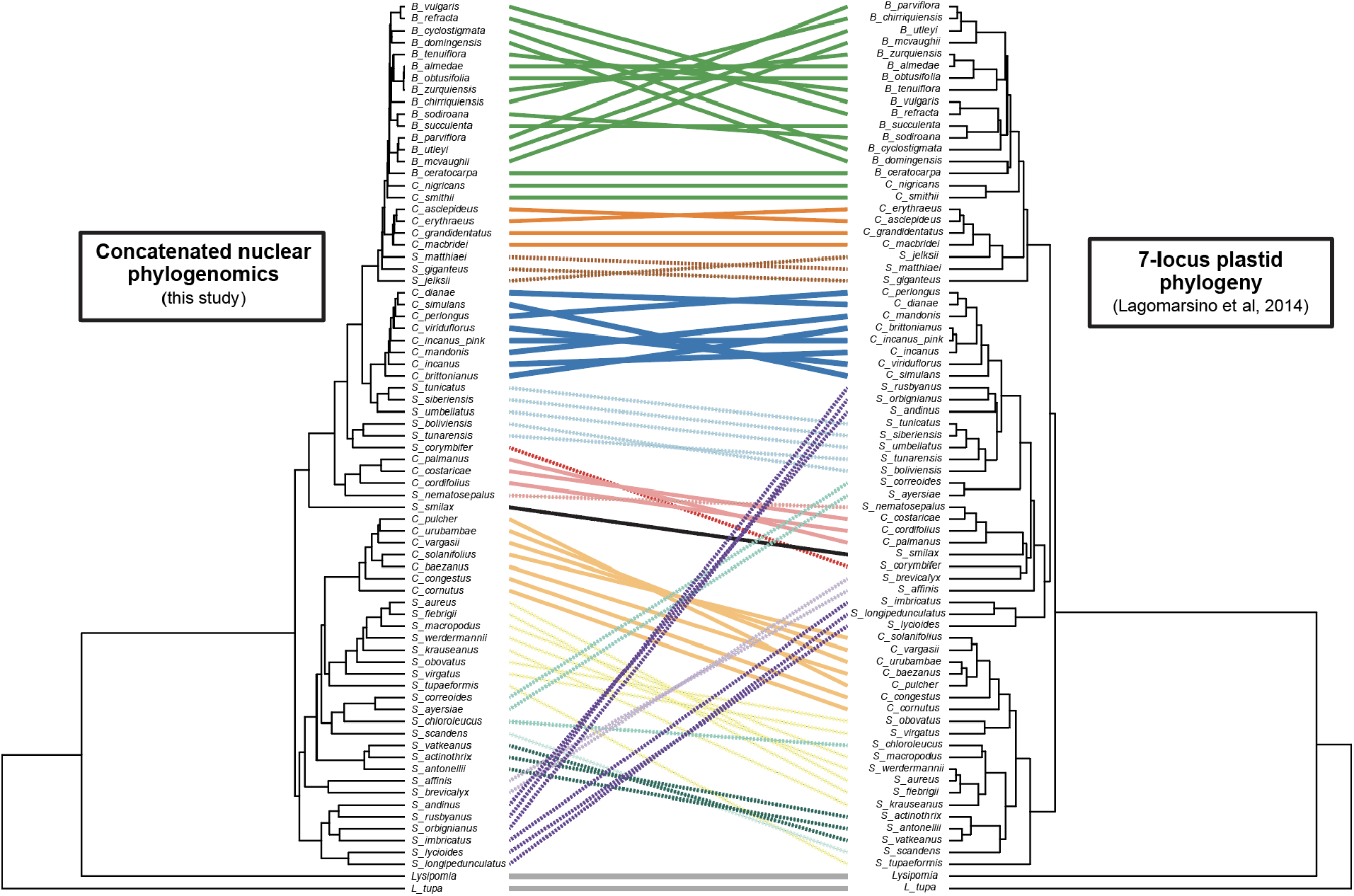
Tanglegram illustrating the conflicts between the RAxML concatenated nuclear phylogenomic topology (left) and the seven locus plastid phylogeny of Lagomarsino et al. (2014; right). Lines are color-coded according to the scheme in Figure 2, with baccate species represented by unbroken lines and capsular species represented by dashed lines.

### Future classification needs of Neotropical Lobelioideae

The classification of Neotropical Lobeloideae is in dire need of a thorough revision. This is especially true for *Siphocampylus*, for which none of the subgeneric taxa of *Siphocampylus* from the latest monograph (Wimmer 1943, 1955, 1968) for which more than a single species was included in our sampling are monophyletic (Fig. 5).

**Figure 5.**
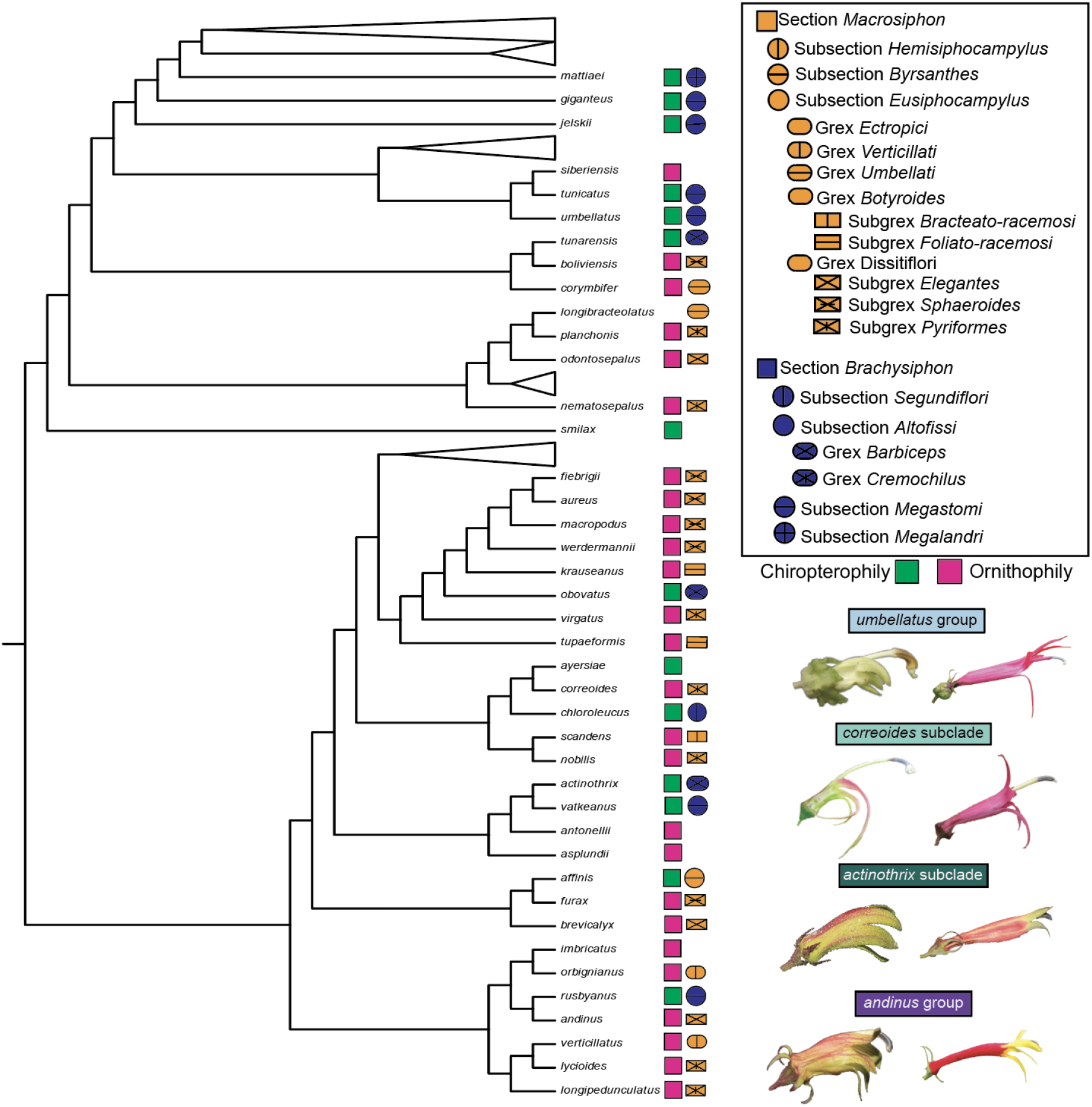
The nonmonophyly that characterizes the current classification *Siphocampylus* is, in part, explained by convergent evolution of floral forms within pollination syndromes as infragenetic groups were circumscribed based on floral traits. Intrageneric classification scheme of Siphocampylus from Wimmer (1968) is presented in legend at top right, with section Macrosiphon in orange and section Brachysiphon in blue, with specific shapes and line patterns delimiting subsectional taxa. The taxa to which Siphocampylus species belong is denoted next to the tip of the RAxML topology, along with whether the species is hummingbird or bat pollinated. Siphocampylus species with no taxonomic symbol next to their name were not included in Wimmer (1968). Examples of closely related pollination shifts shown at bottom right, with chiropterophilous species at left and ornithophilous species at right, with informal clade names above with colors corresponding to Fig. 2. Picture species are *S. tunicatus* and *S. boliviensis* in the *umbellatus* group; *S. ayersiae* and *S. correoides* in the *correoides* group; *S. actinothrix* and *S. antonellii* in the *actinothrix* group; and *S. rusybanus* and *S. orbignianus* in the *andinus* group.

The challenge of reclassification is amplified by the paraphyletic and polyphyletic nature of traits used to define named taxa in this clade. At the genus level, capsular fruits, the defining trait of *Siphocampylus*, are paraphyletic, while the multiple *Centropogon* and *Burmeistera* lineages represent independent evolution of berries (Lagomarsino *et al*. 2014). Similarly, taxa within *Siphocampylus* are largely defined by traits that have undergone repeated convergent evolution. For example, the two sections of *Siphocampylus* largely correspond to hummingbird (section *Macrosiphon*) and bat (section *Brachysiphon*) pollination syndromes (Fig. 5). The former is defined by long, narrow, brightly colored corollas with filaments that are adnate high on the corolla (a potential adaptation to reduce damage from hummingbird bills), while the latter is defined by short, wide, greenish or dull-colored corollas with free filaments or filaments that are adnate near the bear of the corolla. These traits are known to accurately predict pollinators in the centropogonid clade (Muchhala 2003, 2006a; Muchhala and Thomson 2009; Lagomarsino and Muchhala 2019), and undergo strong selection pressure during the frequent shifts between pollination syndromes that characterize the entire centropogonid clade’s history (Lagomarsino *et al*. 2017). This reliance on traits that have been shown to be plesiomorphic (e.g., capsules for *Siphcampylus* and hummingbird pollination for section *Microsiphon*) has resulted in a Russian doll of non-monophyletic classification that will require substantial reconfiguration to accurately reflect evolutionary relationships— a reconfiguration that will require a modification of classification at the level of the Lobelioideae subfamily, which is based on fruit type.

Rapid trait evolution and frequent convergence will make designating new taxa difficult, especially within paraphyletic *Siphocampylus*. There are no obvious floral, fruit, or habit traits separating the various subclades of *Siphocampylus* that we have identified from all other lineages. Instead, traits that may experience less convergent selection pressure, including types of pubescence, aspects of seed morphology or anatomy, and micromorphology, are likely to be more useful. However, even with our substantial taxonomic expertise in this group, no characters are immediately obvious, suggesting that perhaps a combination of these characters may be necessary. Combining morphology and biogeography is likely to most adequately reflect relationships within the clade. Geography has already been shown to be informative: within the clade, the only monophyletic subsection of *Siphocampylus* (subsection *Hemisiphocampylus*), which is distantly related from all other species of the genus, is restricted to the Caribbean, while peruvianid clade of *Centropogon* is restricted to the Central Andes of Peru and Bolivia (Lagomarsino *et al*. 2014).

Finally, determining how to best recircumscribe the three genera comprising the centropogonid clade (i.e., *Centropogon*, *Burmeistera*, and *Siphocampylus*) requires a stable and well-supported phylogeny. While our new phylogeny is an important step in this direction, future phylogenetic research should aim for deeper taxon sampling, with a particularly important goal of increasing sampling of *Siphocampylus* species. Unlike *Centropogon*, for which species can be relatively easily placed into subclades that are unlikely to change with additional taxon sampling, it is possible that we will continue to identify novel lineages of *Siphocampylus* as additional species are added to the phylogeny— such as the *odontosepalus* group in this study, which comprises entirely newly sequenced species. Future reclassification efforts for Neotropical Lobelioideae represent a significant challenge, but one that can be met with additional phylogenetic data, further examination of morphology, and a deep understanding of trait evolution

### Phylogenomic and systematic challenges innate to rapid, morphologically diverse rapid radiations

Difficulty resolving relationships of rapid radiations is a well-known phenomenon in plant systematics, and is exemplified in many Andean plants radiations (Hughes and Eastwood 2006; Nürk *et al*. 2013; Uribe-Convers *et al*. 2016; Pease *et al*. 2016; Vargas *et al*. 2017; Morales-Briones *et al*. 2018; Frost and Lagomarsino 2021). The centropogonid clade, one of the fastest evolutionary radiations reported to date, is no exception. The extensive gene tree discordance we document is expected given that rates of incomplete lineage sorting are inversely proportional with time between speciation events (Pamilo and Nei 1988). Further, the history of this group is likely marked by frequent introgression, as suggested by the cytonuclear discordance and the unstable placement of several rogue species and lineages.

Phylogenomic complexity of the centropogonid clade corresponds with rapid morphological innovation (Lagomarsino *et al*. 2014, 2016, 2017), as has been observed in many distantly related groups of organisms (Parins-Fukuchi *et al*. 2020). It is possible that rampant convergent trait evolution in fruit and floral traits reflects hemiplasy (i.e., traits whose genetic basis is determined by a gene whose history does not match the species tree; (Avise and Robinson 2008; Guerrero and Hahn 2018), rather than de novo convergent evolution. Hemiplasy as the main driver of fruit type evolution seems unlikely in the centropogonid clade, as fruit type is a complex character (Ortiz-Ramírez *et al*. 2018). In the centropogonid clade, baccate fruits tend to be very distinct and readily distinguishable across subclades, suggesting that even if hemiplasy is fully or partially responsible for repeated shifts in fruit type, that the individual genes involved are likely different or there is a degree of de novo variation in addition to historical sorting. However, individual floral traits (including those related to pollination syndrome including color, scent, and overall size) seem more likely to have an extensive history of hemiplasy (Stankowski and Streisfeld 2015). In fact, the paraphyletic phylogenetic distribution of floral traits in the centropogonid clade (vs. the synapomorphic distribution of fruit type) is similar to *Jaltomata*, a tomato relative in which de novo evolution seems to drive fruit traits (in this case, fruit color) and evolution of floral traits likely represents hemiplasy (Wu *et al*. 2018).

Understanding the macroevolutionary history of rapid, species-rich radiations with extensive morphological variation remains an exciting challenge in evolutionary biology. Despite large datasets, support for phylogenomic relationships in these groups tends to be low, likely reflecting the complex, often non-bifurcating lineage splitting events and small populations that characterize these groups. This is true even in well-trodden systems that have the benefit of substantial genomic resources (e.g., Lake Victoria cichlids, (Meier *et al*. 2017); Andean tomatoes, (Pease *et al*. 2016). Andean cloud forest radiations, including the centropogonid clade, exemplify this phylogenomic challenge. It is likely that full understanding of relationships of the understudied, but species-rich plant radiations of this biodiversity hotspot will remain a difficult problem in phylogenomics for years to come, even as datasets continue to expand in size and models improve. This exciting arena of evolutionary biology contrasts with the real threat to biodiversity in this region and ongoing extinction of species that belong to the many rapid radiations whose evolutionary histories we are just beginning to untangle (Antonelli 2021).

## Supporting information

Appendix S1-Voucher Information for all samples included in study.

Appendix S2-Summary statistics for sequence data for all samples included in study.

## Acknowledgements

This research was supported by a National Science Foundation Postdoctoral Research Fellowship in Biology to LPL (Award 1523880), a Field Research for Conservation Grant from the WildCare Institute at the St. Louis Zoo to LPL and NM, and startup funds to LPL from the LSU College of Science and Department of Biological Sciences. AA acknowledges financial support from the Swedish Research Council (2019-05191), the Swedish Foundation for Strategic Research (FFL15-0196) and the Royal Botanic Gardens, Kew.LF was supported by an Open Research Grant from the Grinnell College Center for Careers, Life, and Service. The Muchhala Lab at UMSL, including Serena Achá, Camilo Calderón-Acevedo, Diana Gamba, and Rosana Maguiña, provided assistance in the early stages of this research, and the Lagomarsino Lab at LSU provided feedback on an earlier draft. Brant Faircloth and Carl Oliveros provided assistance in implementing the phyluce pipeline. Portions of this research were conducted with high performance computing resources provided by Louisiana State University (http://www.hpc.lsu.edu). The herbarium at the Missouri Botanical Garden (MO) provided important access to their collections.

## Supplemental Material Legends

**Appendix S1-** Voucher Information for all samples included in study. Tip names corresponding to Sample_ID from Appendix S1.

**Appendix S2-** Summary statistics for sequence data for all samples included in study. Tip names corresponding to Sample_ID from Appendix S1.

**Figure S1.** Results of RAxML analysis without *Burmeistera* collapsed, as in Fig 2. Tip names corresponding to Sample_ID from Appendix S1.

**Figure S2.** Results of ASTRAL analysis without *Burmeistera* collapsed, as in Fig 3a. Tip names corresponding to Sample_ID from Appendix S1.

**Figure S3.** Results of ASTRID analysis without *Burmeistera* collapsed, as in Fig 3b. Tip names corresponding to Sample_ID from Appendix S1.

**Figure S4.** Results of SVDquartets analysis without *Burmeistera* collapsed, as in Fig 3c. Tip names corresponding to Sample_ID from Appendix S1.

**Figure S5.** Results of Phyparts analysis along RAxML phylogeny, including discordance visualizations for every node in phylogeny. Colors in pie chart correspond to the proportion of gene trees that fall into different categories of concordance (blue: concordant genes; green: most common conflicting bipartition; red: other conflicting bipartitions; grey: gene trees with no information). Tip names corresponding to Sample_ID from Appendix S1.

## Literature Cited

1. Acha S, Linan A, MacDougal J, Edwards C. 2021. The evolutionary history of vines in a neotropical biodiversity hotspot: Phylogenomics and biogeography of a large passion flower clade (*Passiflora* section *Decaloba*). Molecular phylogenetics and evolution 164: 107260.

2. Alsos IG, Lavergne S, Merkel MKF, et al. 2020. The treasure vault can be opened: Large-scale genome skimming works well using herbarium and silica gel dried material. Plants 9.

3. Antonelli A. 2008. Higher level phylogeny and evolutionary trends in Campanulaceae subfam. Lobelioideae: molecular signal overshadows morphology. Molecular phylogenetics and evolution 46: 1–18.

4. Antonelli A. 2009. Have giant lobelias evolved several times independently? Life form shifts and historical biogeography of the cosmopolitan and highly diverse subfamily Lobelioideae (Campanulaceae). BMC Biology 7: 82.

5. Antonelli A. 2021. The rise and fall of Neotropical biodiversity. Botanical journal of the Linnean Society. Linnean Society of London: https://doi.org/10.1093/botlinnean/boab061.

6. Avise JC, Robinson TJ. 2008. Hemiplasy: a new term in the lexicon of phylogenetics. Systematic biology 57: 503–507.

7. Bagley JC, Uribe-Convers S, Carlsen MM, Muchhala N. 2020. Utility of targeted sequence capture for phylogenomics in rapid, recent angiosperm radiations: Neotropical *Burmeistera* bellflowers as a case study. Molecular phylogenetics and evolution: 106769.

8. Bakker FT. 2017. Herbarium genomics: skimming and plastomics from archival specimens. Webbia 72: 35–45.

9. Boehm MM, Scholer MN, Kennedy JJ, et al. 2018. The Manú Gradient as a study system for bird pollination. Biodiversity data journal: e22241.

10. Bolger AM, Lohse M, Usadel B. 2014. Trimmomatic: a flexible trimmer for Illumina sequence data. Bioinformatics 30: 2114–2120.

11. Bravo GA, Antonelli A, Bacon CD, et al. 2019. Embracing heterogeneity: coalescing the Tree of Life and the future of phylogenomics. PeerJ 7: e6399.

12. Brewer GE, Clarkson JJ, Maurin O, et al. 2019. Factors affecting targeted sequencing of 353 nuclear genes From herbarium specimens spanning the diversity of angiosperms. Frontiers in plant science 10: 1102.

13. Castresana J. 2000. Selection of conserved blocks from multiple alignments for their use in phylogenetic analysis. Molecular biology and evolution 17: 540–552.

14. Chifman J, Kubatko L. 2014. Quartet inference from SNP data under the coalescent model. Bioinformatics 30: 3317–3324.

15. Crowl AA, Miles NW, Visger CJ, et al. 2016. A global perspective on Campanulaceae: Biogeographic, genomic, and floral evolution. American journal of botany 103: 233–245.

16. Doyle JJ. 2021. Defining coalescent genes: Theory meets practice in organelle phylogenomics. Systematic biology: https://doi.org/10.1093/sysbio/syab053.

17. Doyle JJ, Doyle JL. 1987. CTAB DNA extraction in plants. Phytochemical Bulletin 19: 11–15.

18. Edwards SV. 2009. Is a new and general theory of molecular systematics emerging? Evolution; international journal of organic evolution 63: 1–19.

19. Faircloth BC. 2013. Illumiprocessor: a trimmomatic wrapper for parallel adapter and quality trimming. *See* http://dx.doi.org/10.6079/J9ILL *(accessed 4 November 2016)*.

20. Faircloth BC. 2016. PHYLUCE is a software package for the analysis of conserved genomic loci. Bioinformatics 32: 786–788.

21. Faircloth BC, McCormack JE, Crawford NG, Harvey MG, Brumfield RT, Glenn TC. 2012. Ultraconserved elements anchor thousands of genetic markers spanning multiple evolutionary timescales. Systematic biology 61: 717–726.

22. Firneno TJ Jr, O’Neill JR, Portik DM, Emery AH, Townsend JH, Fujita MK. 2020. Finding complexity in complexes: Assessing the causes of mitonuclear discordance in a problematic species complex of Mesoamerican toads. Molecular ecology 29: 3543–3559.

23. Frost L, Lagomarsino L. 2021. More-curated data outperforms more data: Treatment of cryptic and known paralogs improves phylogenomic analysis and resolves a northern Andean origin of Freziera (Pentaphylacaceae). *bioRxiv*: 2021.07.01.450750.

24. Galili T. 2015. dendextend: an R package for visualizing, adjusting and comparing trees of hierarchical clustering. Bioinformatics 31: 3718–3720.

25. Gonçalves DJP, Simpson BB, Ortiz EM, Shimizu GH, Jansen RK. 2019. Incongruence between gene trees and species trees and phylogenetic signal variation in plastid genes. Molecular phylogenetics and evolution 138: 219–232.

26. Grabherr MG, Haas BJ, Yassour M, et al. 2011. Full-length transcriptome assembly from RNA-Seq data without a reference genome. Nature biotechnology 29: 644–652.

27. Guerrero RF, Hahn MW. 2018. Quantifying the risk of hemiplasy in phylogenetic inference. Proceedings of the National Academy of Sciences of the United States of America 115: 12787–12792.

28. Hart ML, Forrest LL, Nicholls JA, Kidner CA. 2016. Retrieval of hundreds of nuclear loci from herbarium specimens. Taxon 65: 1081–1092.

29. Hosner PA, Faircloth BC, Glenn TC, Braun EL, Kimball RT. 2016. Avoiding missing data biases in phylogenomic inference: An empirical study in the landfowl (Aves: Galliformes). Molecular biology and evolution 33: 1110–1125.

30. Hughes CE, Eastwood R. 2006. Island radiation on a continental scale: exceptional rates of plant diversification after uplift of the Andes. Proceedings of the National Academy of Sciences of the United States of America 103: 10334–10339.

31. Johnson MG, Gardner EM, Liu Y, et al. 2016. HybPiper: Extracting coding sequence and introns for phylogenetics from high-throughput sequencing reads using target enrichment. Applications in plant sciences 4: 1600016.

32. Jones G, Sagitov S, Oxelman B. 2013. Statistical inference of allopolyploid species networks in the presence of incomplete lineage sorting. Systematic biology 62: 467–478.

33. Kates HR, Doby JR, Siniscalchi CM, et al. 2021. The effects of herbarium specimen characteristics on short-read NGS sequencing success in nearly 8000 specimens: Old, degraded samples have lower DNA yields but consistent sequencing success. Frontiers in plant science 12: 669064.

34. Katoh K, Standley DM. 2013. MAFFT multiple sequence alignment software version 7: Improvements in performance and usability. Molecular biology and evolution 30: 772–780.

35. Knox EB, Muasya AM, Muchhala N. 2008. The predominantly South American clade of Lobeliaceae. Systematic botany 33: 462–468.

36. Lagomarsino LP, Antonelli A, Muchhala N, Timmermann A, Mathews S, Davis CC. 2014. Phylogeny, classification, and fruit evolution of the species-rich Neotropical bellflowers (Campanulaceae: Lobelioideae). American journal of botany 101: 2097–2112.

37. Lagomarsino LP, Condamine FL, Antonelli A, Mulch A, Davis CC. 2016. The abiotic and biotic drivers of rapid diversification in Andean bellflowers (Campanulaceae). The New phytologist 210: 1430–1442.

38. Lagomarsino LP, Forrestel EJ, Muchhala N, Davis CC. 2017. Repeated evolution of vertebrate pollination syndromes in a recently diverged Andean plant clade. Evolution; international journal of organic evolution 71: 1970–1985.

39. Lagomarsino LP, Muchhala N. 2019. A gradient of pollination specialization in three species of Bolivian Centropogon. American journal of botany 106: 633–642.

40. Lagomarsino LP, Santamaría-Aguilar D. 2016. Two new species of *Siphocampylus* (Campanulaceae, Lobelioideae) from the Central Andes. PhytoKeys: 105–117.

41. Lammers TG. 1998. Review of the Neotropical Endemics Burmeistera, Centropogon, and Siphocampylus (Campanulaceae: Lobelioideae), with Description of 18 New Species and a New Section. Brittonia 50: 233–262.

42. Lanfear R, Frandsen PB, Wright AM, Senfeld T, Calcott B. 2017. PartitionFinder 2: New methods for selecting partitioned models of evolution for molecular and morphological phylogenetic analyses. Molecular biology and evolution 34: 772–773.

43. Lee-Yaw JA, Grassa CJ, Joly S, Andrew RL, Rieseberg LH. 2019. An evaluation of alternative explanations for widespread cytonuclear discordance in annual sunflowers (*Helianthus*). The New phytologist 221: 515–526.

44. Lemmon AR, Emme SA, Lemmon EM. 2012. Anchored hybrid enrichment for massively high-throughput phylogenomics. Systematic biology 61: 727–744.

45. Lendemer J, Thiers B, Monfils AK, et al. 2020. The Extended Specimen Network: A strategy to enhance US biodiversity collections, promote research and education. Bioscience 70: 23–30.

46. McKain MR, Johnson MG, Uribe-Convers S, Eaton D, Yang Y. 2018. Practical considerations for plant phylogenomics. Applications in Plant Sciences 6: e1038.

47. McVaugh R. 1965. South American Lobelioideae new to science. Annals of the Missouri Botanical Garden. Missouri Botanical Garden 52: 399.

48. Meier JI, Marques DA, Mwaiko S, Wagner CE, Excoffier L, Seehausen O. 2017. Ancient hybridization fuels rapid cichlid fish adaptive radiations. Nature communications 8: 14363.

49. Molloy EK, Warnow T. 2018. To include or not to include: The impact of gene filtering on species tree estimation methods. Systematic biology 67: 285–303.

50. Morales-Briones DF, Liston A, Tank DC. 2018. Phylogenomic analyses reveal a deep history of hybridization and polyploidy in the Neotropical genus *Lachemilla* (Rosaceae). The New phytologist 218: 1668–1684.

51. Muchhala N. 2003. Exploring the boundary between pollination syndromes: Bats and hummingbirds as pollinators of *Burmeistera cyclostigmata* and *B. tenuiflora* (Campanulaceae). Oecologia 134: 373–380.

52. Muchhala N. 2006a. The pollination biology of *Burmeistera* (Campanulaceae): Specialization and syndromes. American journal of botany 93: 1081–1089.

53. Muchhala N. 2006b. Nectar bat stows huge tongue in its rib cage. Nature 444: 701–702.

54. Muchhala N, Potts MD. 2007. Character displacement among bat-pollinated flowers of the genus *Burmeistera*: Analysis of mechanism, process and pattern. Proceedings of the Royal Society B: Biological Sciences 274: 2731–2737.

55. Muchhala N, Thomson JD. 2009. Going to great lengths: selection for long corolla tubes in an extremely specialized bat-flower mutualism. Proceedings. Biological sciences / The Royal Society 276: 2147–2152.

56. Nevado B, Atchison GW, Hughes CE, Filatov DA. 2016. Widespread adaptive evolution during repeated evolutionary radiations in New World lupins. Nature communications 7: 12384.

57. Nürk NM, Scheriau C, Madriñán S. 2013. Explosive radiation in high Andean *Hypericum*—rates of diversification among New World lineages. Frontiers in Genetics 4: 175.

58. Ogilvie HA, Bouckaert RR, Drummond AJ. 2017. StarBEAST2 brings faster species tree inference and accurate estimates of substitution rates. Molecular biology and evolution 34: 2101–2114.

59. Ortiz-Ramírez CI, Plata-Arboleda S, Pabón-Mora N. 2018. Evolution of genes associated with gynoecium patterning and fruit development in Solanaceae. Annals of botany 121: 1211–1230.

60. Pamilo P, Nei M. 1988. Relationships between gene trees and species trees. Molecular biology and evolution 5: 568–583.

61. Parins-Fukuchi C, Stull GW, Smith SA. 2020. Phylogenomic conflict coincides with rapid morphological innovation. Cold Spring Harbor Laboratory: 2020.11.04.368902.

62. Pease JB, Haak DC, Hahn MW, Moyle LC. 2016. Phylogenomics reveals three sources of adaptive variation during a rapid radiation. PLoS biology 14: e1002379.

63. Pouchon C, Fernández A, Nassar JM, et al. 2018. Phylogenomic Analysis of the Explosive Adaptive Radiation of the *Espeletia* Complex (Asteraceae) in the Tropical Andes. Systematic biology 67: 1041–1060.

64. Rabeler RK, Svoboda HT, Thiers B, et al. 2019. Herbarium practices and ethics, III. Systematic Botany 44: 7–13.

65. Reiseberg L, Soltis D. 1991. Phylogenetic consequences of cytoplasmic gene flow in plants. Evolutionary Trends in Plants 5: 65–84.

66. Rothfels CJ. 2021. Polyploid phylogenetics. The New phytologist 230: 66–72.

67. Schneider JV, Jungcurt T, Cardoso D, et al. 2021. Phylogenomics of the tropical plant family Ochnaceae using targeted enrichment of nuclear genes and 250+ taxa. Taxon 70: 48–71.

68. Silva C, Besnard G, Piot A, Razanatsoa J, Oliveira RP, Vorontsova MS. 2017. Museomics resolve the systematics of an endangered grass lineage endemic to north-western Madagascar. Annals of botany 119: 339–351.

69. Sloan DB, Havird JC, Sharbrough J. 2017. The on-again, off-again relationship between mitochondrial genomes and species boundaries. Molecular ecology 26: 2212–2236.

70. Smith SA, Moore MJ, Brown JW, Yang Y. 2015. Analysis of phylogenomic datasets reveals conflict, concordance, and gene duplications with examples from animals and plants. BMC evolutionary biology 15: 150.

71. Solís-Lemus C, Bastide P, Ané C. 2017. PhyloNetworks: A package for phylogenetic networks. Molecular biology and evolution 34: 3292–3298.

72. Stankowski S, Streisfeld MA. 2015. Introgressive hybridization facilitates adaptive divergence in a recent radiation of monkeyflowers. Proceedings. Biological sciences / The Royal Society 282: 20151666.

73. Stein BA. 1987. Systematics and evolution of Centropogon subgenus Centropogon (Campanulaceae: Lobelioideae) (PH Raven, Ed.).

74. Stein BA. 1992. Sicklebill hummingbirds, ants, and flowers: Plant-animal interactions and evolutionary relationships in Andean Lobeliaceae. Bioscience 42: 27–33.

75. Swofford DL. 2001. PAUP*: Phylogenetic Analysis Using Parsimony (and other methods) 4.0.b5.

76. Than C, Ruths D, Nakhleh L. 2008. PhyloNet: a software package for analyzing and reconstructing reticulate evolutionary relationships. BMC bioinformatics 9: 322.

77. Thomas AE, Igea J, Meudt HM, Albach DC, Lee WG, Tanentzap AJ. 2021. Using target sequence capture to improve the phylogenetic resolution of a rapid radiation in New Zealand *Veronica*. American journal of botany 108: 1289–1306.

78. Uribe-Convers S, Settles ML, Tank DC. 2016. A phylogenomic approach based on PCR target enrichment and high throughput sequencing: Resolving the diversity within the South American Species of *Bartsia* L. (Orobanchaceae). PLOS ONE 11: e0148203.

79. Vachaspati P, Warnow T. 2015. ASTRID: Accurate Species TRees from Internode Distances. BMC genomics 16: S3.

80. Vargas OM, Ortiz EM, Simpson BB. 2017. Conflicting phylogenomic signals reveal a pattern of reticulate evolution in a recent high-Andean diversification (Asteraceae: Astereae: *Diplostephium*). The New phytologist 214: 1736–1750.

81. Weitemier K, Straub SCK, Cronn RC, et al. 2014. Hyb-Seq: Combining target enrichment and genome skimming for plant phylogenomics. Applications in plant sciences 2: 1400042.

82. Wimmer FE. 1943. Campanulaceae-Lobelioideae. I. Teil In: Mansfeld R, ed. Das Pflanzenreich IV.276b. Leipzig: Wilhem Engelmann, 1–260.

83. Wimmer FE. 1955. Lobeliacearum species Novae Austro-Americanae. Brittonia 8: 107–111.

84. Wimmer FE. 1968. Campanulaceae–Lobelioideae supplementum et Campanulaceae–Cyphiodeae In: Danert S, ed. Das Pflanzenreich. Berlin, Germany: Akademie-Verlag, 815–1024.

85. Wu M, Kostyun JL, Hahn MW, Moyle LC. 2018. Dissecting the basis of novel trait evolution in a radiation with widespread phylogenetic discordance. Molecular ecology: 10.1111/mec.14780.

86. Zhang C, Sayyari E, Mirarab S. 2017. ASTRAL-III: Increased scalability and impacts of contracting low support branches In: Lecture Notes in Computer Science (including subseries Lecture Notes in Artificial Intelligence and Lecture Notes in Bioinformatics). Springer, Cham, 53–75.

