## Supplemental Figures for "Increased resolution in the face of conflict: phylogenomics of the Neotropical bellflowers (Campanulaceae: Lobelioideae), a rapid plant radiation"

**Figure S1.** Results of RAxML analysis without Burmeistera collapsed, as in Fig 2. Tip names corresponding to Sample_ID from Appendix S1.

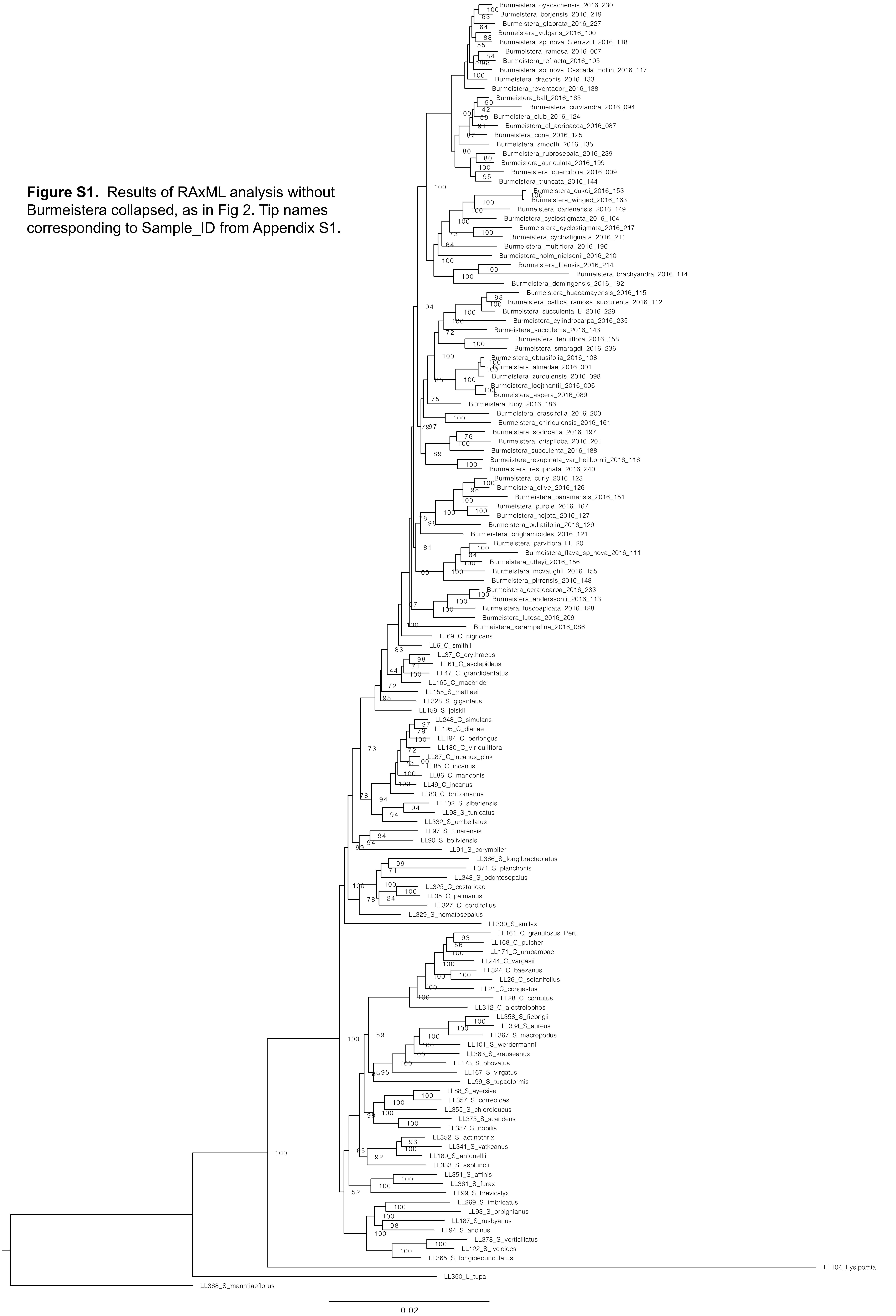

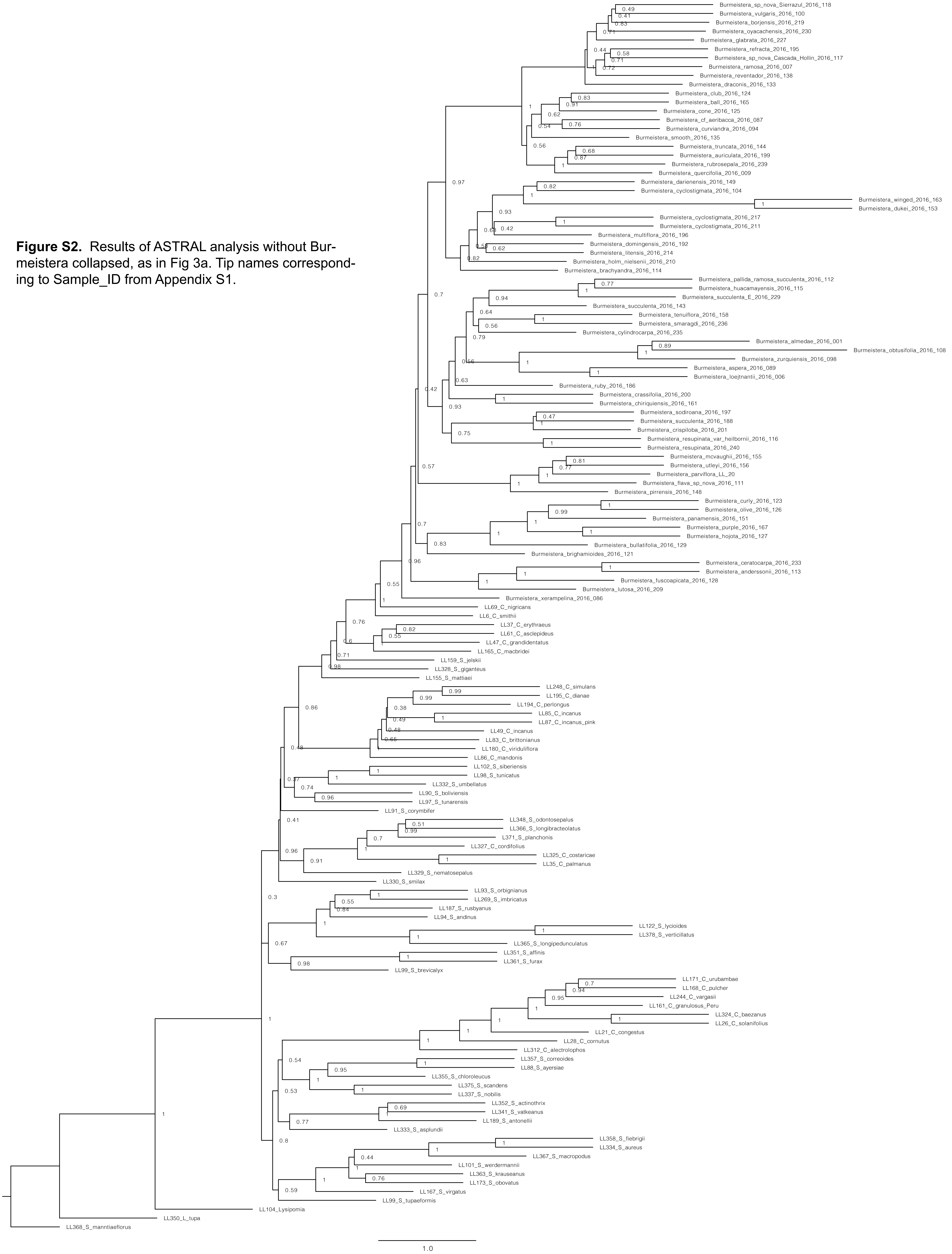


**Figure S2.** Results of ASTRAL analysis without Burmeistera collapsed, as in Fig 3a. Tip names corresponding to Sample_ID from Appendix S1.

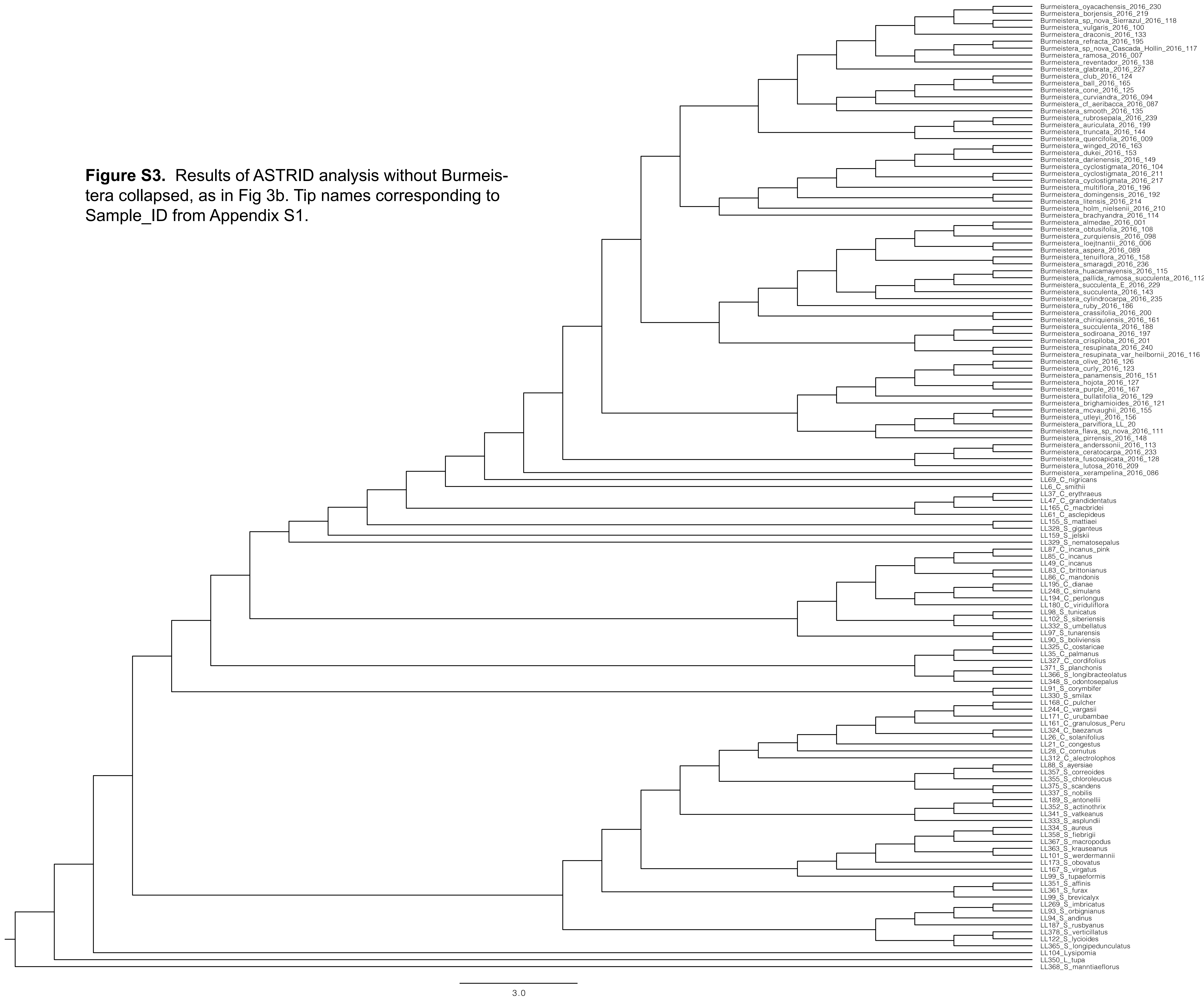


**Figure S3.** Results of ASTRID analysis without Burmeistera collapsed, as in Fig 3b. Tip names corresponding to Sample_ID from Appendix S1.

**Figure S4.** Results of SVDquartets analysis without Burmeistera collapsed, as in Fig 3c. Tip names corresponding to Sample_ID from Appendix S1.

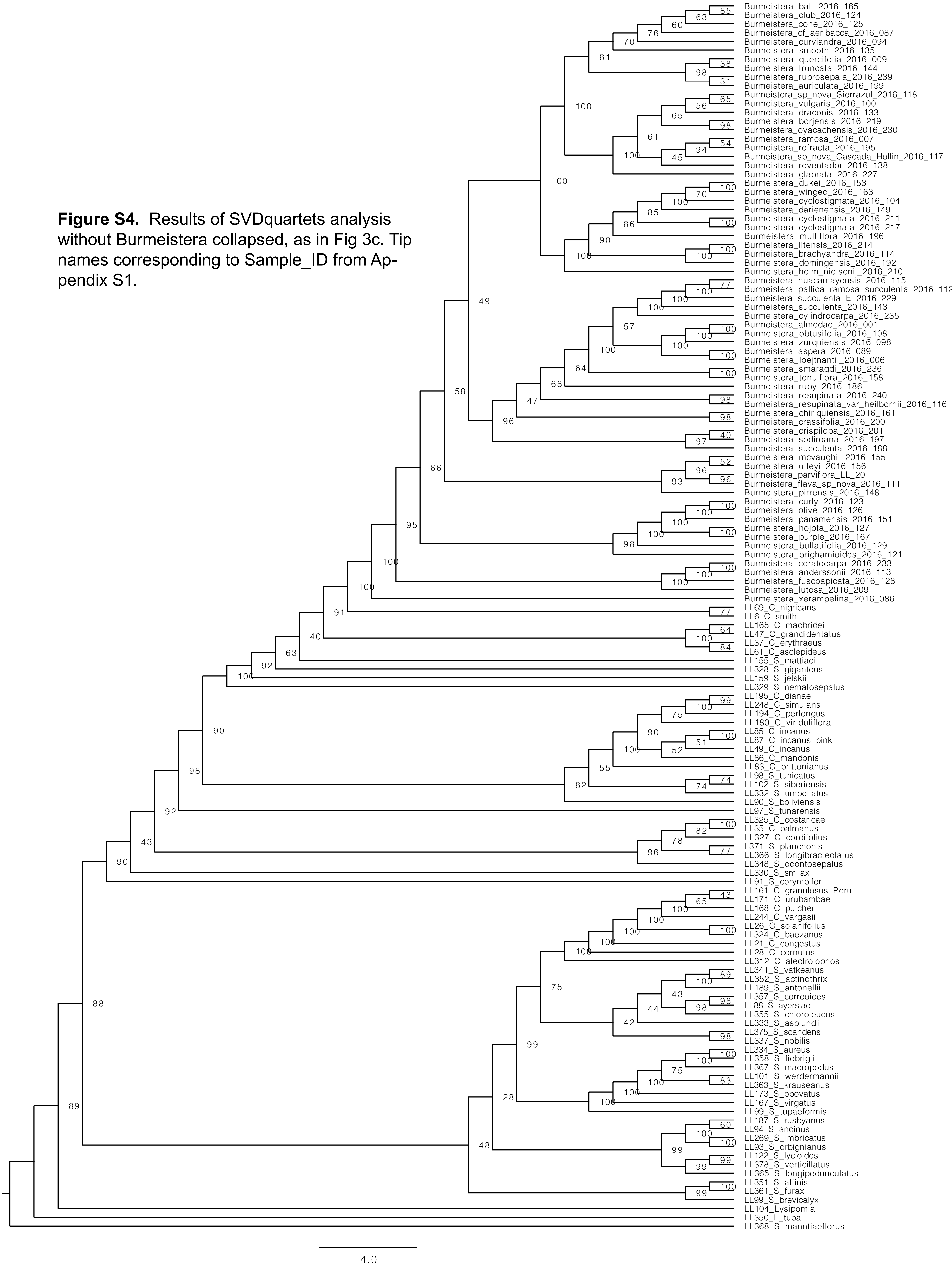

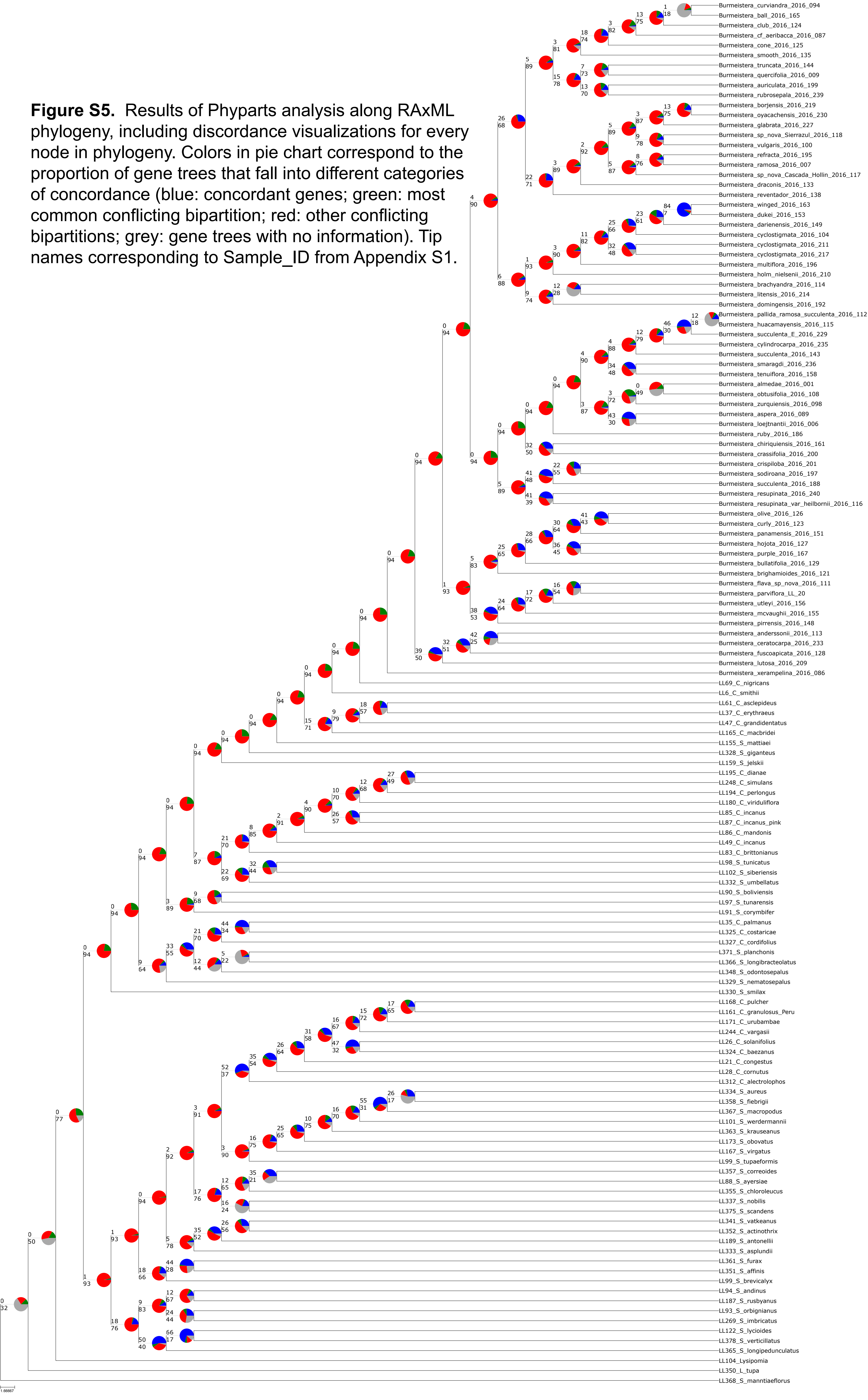


**Figure S5.** Results of Phyparts analysis along RAxML phylogeny, including discordance visualizations for every node in phylogeny. Colors in pie chart correspond to the proportion of gene trees that fall into different categories of concordance (blue: concordant genes; green: most common conflicting bipartition; red: other conflicting bipartitions; grey: gene trees with no information). Tip names corresponding to Sample_ID from Appendix S1.
